# Modern Hopfield Networks and Attention for Immune Repertoire Classification

**DOI:** 10.1101/2020.04.12.038158

**Authors:** Michael Widrich, Bernhard Schäfl, Milena Pavlović, Hubert Ramsauer, Lukas Gruber, Markus Holzleitner, Johannes Brandstetter, Geir Kjetil Sandve, Victor Greiff, Sepp Hochreiter, Günter Klambauer

## Abstract

A central mechanism in machine learning is to identify, store, and recognize patterns. How to learn, access, and retrieve such patterns is crucial in Hopfield networks and the more recent transformer architectures. We show that the attention mechanism of transformer architectures is actually the update rule of modern Hopfield networks that can store exponentially many patterns. We exploit this high storage capacity of modern Hopfield networks to solve a challenging multiple instance learning (MIL) problem in computational biology: immune repertoire classification. Accurate and interpretable machine learning methods solving this problem could pave the way towards new vaccines and therapies, which is currently a very relevant research topic intensified by the COVID-19 crisis. Immune repertoire classification based on the vast number of immunosequences of an individual is a MIL problem with an unprecedentedly massive number of instances, two orders of magnitude larger than currently considered problems, and with an extremely low witness rate. In this work, we present our novel method DeepRC that integrates transformer-like attention, or equivalently modern Hopfield networks, into deep learning architectures for massive MIL such as immune repertoire classification. We demonstrate that DeepRC outperforms all other methods with respect to predictive performance on large-scale experiments, including simulated and real-world virus infection data, and enables the extraction of sequence motifs that are connected to a given disease class. Source code and datasets: *https://github.com/ml-jku/DeepRC*

## Introduction

Transformer architectures (Vaswani et al., 2017) and their attention mechanisms are currently used in many applications, such as natural language processing (NLP), imaging, and also in multiple instance learning (MIL) problems (Lee et al., 2019). In MIL, a set or bag of objects is labelled rather than objects themselves as in standard supervised learning tasks (Dietterich et al., 1997). Examples for MIL problems are medical images, in which each sub-region of the image represents an instance, video classification, in which each frame is an instance, text classification, where words or sentences are instances of a text, point sets, where each point is an instance of a 3D object, and remote sensing data, where each sensor is an instance (Carbonneau et al., 2018; Uriot, 2019). Attention-based MIL has been successfully used for image data, for example to identify tiny objects in large images (Ilse et al., 2018; Pawlowski et al., 2019; Tomita et al., 2019; Kimeswenger et al., 2019) and transformer-like attention mechanisms for sets of points and images (Lee et al., 2019).

However, in MIL problems considered by machine learning methods up to now, the number of instances per bag is in the range of hundreds or few thousands (Carbonneau et al., 2018; Lee et al., 2019) (see also Tab. A2). At the same time the witness rate (WR), the rate of discriminating instances per bag, is already considered low at 1% − 5%. We will tackle the problem of *immune repertoire classification* with hundreds of thousands of instances per bag without instance-level labels and with extremely low witness rates down to 0.01% using an attention mechanism. We show that the attention mechanism of transformers is the update rule of modern Hopfield networks (Krotov & Hopfield, 2016, 2018; Demircigil et al., 2017) that are generalized to continuous states in contrast to classical Hopfield networks (Hopfield, 1982). A detailed derivation and analysis of modern Hopfield networks is given in our companion paper (Ramsauer et al., 2020). These novel continuous state Hopfield networks allow to store and retrieve exponentially (in the dimension of the space) many patterns (see next Section). Thus, modern Hopfield networks with their update rule, which are used as an attention mechanism in the transformer, enable immune repertoire classification in computational biology.

Immune repertoire classification, i.e. classifying the immune status based on the immune repertoire sequences, is essentially a text-book example for a *multiple instance learning* problem (Dietterich et al., 1997; Maron & Lozano-Pérez, 1998; Wang et al., 2018). Briefly, the immune repertoire of an individual consists of an immensely large bag of immune receptors, represented as amino acid sequences. Usually, the presence of only a small fraction of particular receptors determines the immune status with respect to a particular disease (Christophersen et al., 2014; Emerson et al., 2017). This is because the immune system has already acquired a resistance if one or few particular immune receptors that can bind to the disease agent are present. Therefore, classification of immune repertoires bears a high difficulty since each immune repertoire can contain millions of sequences as instances with only a few indicating the class. Further properties of the data that complicate the problem are: (a) The overlap of immune repertoires of different individuals is low (in most cases, maximally low single-digit percentage values) (Greiff et al., 2017; Elhanati et al., 2018), (b) multiple different sequences can bind to the same pathogen (Wucherpfennig et al., 2007), and (c) only subsequences within the sequences determine whether binding to a pathogen is possible (Dash et al., 2017; Glanville et al., 2017; Akbar et al., 2019; Springer et al., 2020; Fischer et al., 2019). In summary, immune repertoire classification can be formulated as multiple instance learning with an extremely low witness rate and large numbers of instances, which represents a challenge for currently available machine learning methods. Furthermore, the methods should ideally be interpretable, since the extraction of class-associated sequence motifs is desired to gain crucial biological insights.

The acquisition of human immune repertoires has been enabled by immunosequencing technology (Georgiou et al., 2014; Brown et al., 2019) which allows to obtain the immune receptor sequences and immune repertoires of individuals. Each individual is uniquely characterized by their immune repertoire, which is acquired and changed during life. This repertoire may be influenced by all diseases that an individual is exposed to during their lives and hence contains highly valuable information about those diseases and the individual’s immune status. Immune receptors enable the immune system to specifically recognize disease agents or pathogens. Each immune encounter is recorded as an immune event into immune memory by preserving and amplifying immune receptors in the repertoire used to fight a given disease. This is, for example, the working principle of vaccination. Each human has about 10^7^–10^8^ unique immune receptors with low overlap across individuals and sampled from a potential diversity of *>* 10^14^ receptors (Mora & Walczak, 2019). The ability to sequence and analyze human immune receptors at large scale has led to fundamental and mechanistic insights into the adaptive immune system and has also opened the opportunity for the development of novel diagnostics and therapy approaches (Georgiou et al., 2014; Brown et al., 2019).

Immunosequencing data have been analyzed with computational methods for a variety of different tasks (Greiff et al., 2015; Shugay et al., 2015; Miho et al., 2018; Yaari & Kleinstein, 2015; Wardemann & Busse, 2017). A large part of the available machine learning methods for immune receptor data has been focusing on the individual immune receptors in a repertoire, with the aim to, for example, predict the antigen or antigen portion (epitope) to which these sequences bind or to predict sharing of receptors across individuals (Gielis et al., 2019; Springer et al., 2020; Jurtz et al., 2018; Moris et al., 2019; Fischer et al., 2019; Greiff et al., 2017; Sidhom et al., 2019; Elhanati et al., 2018). Recently, Jurtz et al. (2018) used 1D convolutional neural networks (CNNs) to predict antigen binding of T-cell receptor (TCR) sequences (specifically, binding of TCR sequences to peptide-MHC complexes) and demonstrated that motifs can be extracted from these models. Similarly, Konishi et al. (2019) use CNNs, gradient boosting, and other machine learning techniques on B-cell receptor (BCR) sequences to distinguish tumor tissue from normal tissue. However, the methods presented so far predict a particular class, the epitope, based on a single input sequence.

Immune repertoire classification has been considered as a MIL problem in the following publications. A Deep Learning framework called DeepTCR (Sidhom et al., 2019) implements several Deep Learning approaches for immunosequencing data. The computational framework, inter alia, allows for attention-based MIL repertoire classifiers and implements a basic form of attention-based averaging. Ostmeyer et al. (2019) already suggested a MIL method for immune repertoire classification. This method considers 4-mers, fixed sub-sequences of length 4, as instances of an input object and trained a logistic regression model with these 4-mers as input. The predictions of the logistic regression model for each 4-mer were max-pooled to obtain one prediction per input object. This approach is characterized by (a) the rigidity of the k-mer features as compared to convolutional kernels (Alipanahi et al., 2015; Zhou & Troyanskaya, 2015; Zeng et al., 2016), (b) the max-pooling operation, which constrains the network to learn from a single, top-ranked k-mer for each iteration over the input object, and (c) the pooling of prediction scores rather than representations (Wang et al., 2018). Our experiments also support that these choices in the design of the method can lead to constraints on the predictive performance (see Table 1).

**Table 1:**
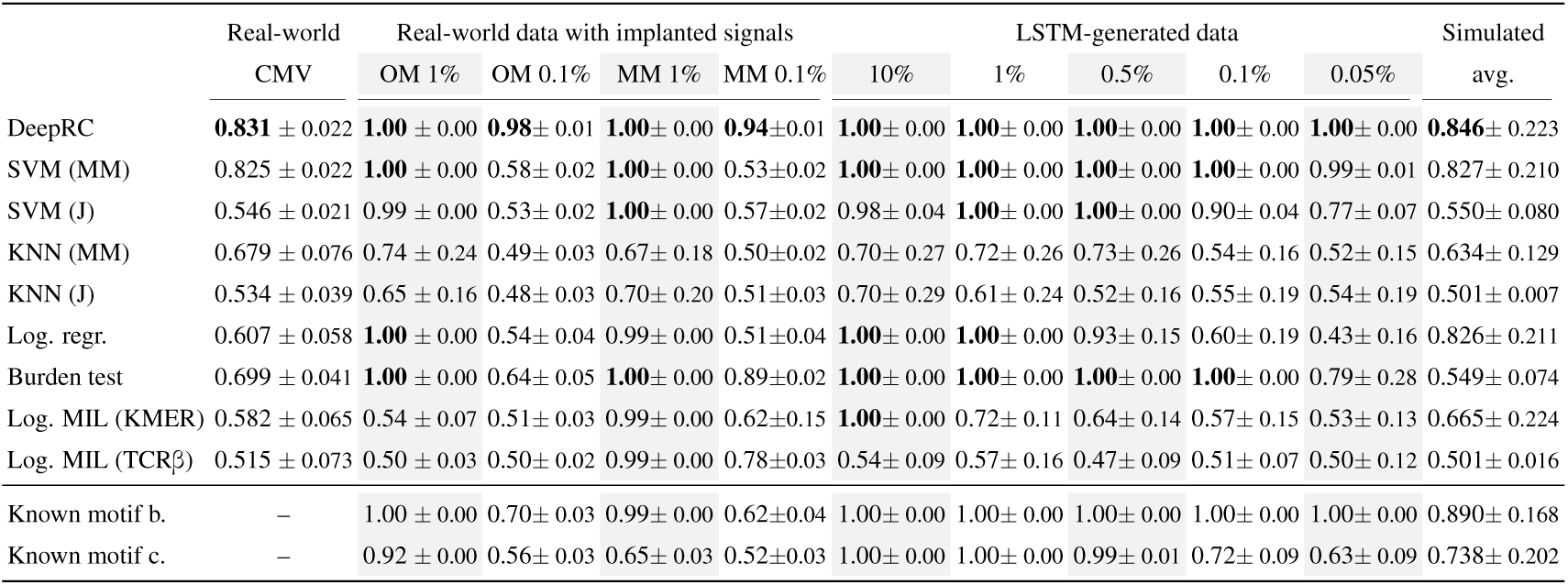
Results in terms of AUC of the competing methods on all datasets. The reported errors are standard deviations across 5 cross-validation (CV) folds (except for the column “Simulated”). **Real-world CMV:** Average performance over 5 CV folds on the *CMV dataset* (Emerson et al., 2017). **Real-world data with implanted signals:** Average performance over 5 CV folds for each of the four datasets. A signal was implanted with a frequency (=witness rate) of 1% or 0.1%. Either a single motif (“OM”) or multiple motifs (“MM”) were implanted. **LSTM-generated data:** Average performance over 5 CV folds for each of the 5 datasets. In each dataset, a signal was implanted with a frequency of 10%, 1%, 0.5%, 0.1%, or 0.05%, respectively. **Simulated:** Here we report the mean over 18 simulated datasets with implanted signals and varying difficulties (see Tab. A9 for details). The error reported is the standard deviation of the AUC values across the 18 datasets.

|  | Real-world | Real-world data with implanted signals |  |  |  | LSTM-generated data |  |  |  |  | Simulated |
| --- | --- | --- | --- | --- | --- | --- | --- | --- | --- | --- | --- |
|  | CMV | OM 1% | OM 0.1% | MM 1% | MM 0.1% | 10% | 1% | 0.5% | 0.1% | 0.05% | avg. |
| DeepRC | $0.831 \pm 0.022$ | $1.00 \pm 0.00$ | $0.98 \pm 0.01$ | $1.00 \pm 0.00$ | $0.94 \pm 0.01$ | $1.00 \pm 0.00$ | $1.00 \pm 0.00$ | $1.00 \pm 0.00$ | $1.00 \pm 0.00$ | $1.00 \pm 0.00$ | $0.846 \pm 0.223$ |
| SVM (MM) | $0.825 \pm 0.022$ | $1.00 \pm 0.00$ | $0.58 \pm 0.02$ | $1.00 \pm 0.00$ | $0.53 \pm 0.02$ | $1.00 \pm 0.00$ | $1.00 \pm 0.00$ | $1.00 \pm 0.00$ | $1.00 \pm 0.00$ | $0.99 \pm 0.01$ | $0.827 \pm 0.210$ |
| SVM (J) | $0.546 \pm 0.021$ | $0.99 \pm 0.00$ | $0.53 \pm 0.02$ | $1.00 \pm 0.00$ | $0.57 \pm 0.02$ | $0.98 \pm 0.04$ | $1.00 \pm 0.00$ | $1.00 \pm 0.00$ | $0.90 \pm 0.04$ | $0.77 \pm 0.07$ | $0.550 \pm 0.080$ |
| KNN (MM) | $0.679 \pm 0.076$ | $0.74 \pm 0.24$ | $0.49 \pm 0.03$ | $0.67 \pm 0.18$ | $0.50 \pm 0.02$ | $0.70 \pm 0.27$ | $0.72 \pm 0.26$ | $0.73 \pm 0.26$ | $0.54 \pm 0.16$ | $0.52 \pm 0.15$ | $0.634 \pm 0.129$ |
| KNN (J) | $0.534 \pm 0.039$ | $0.65 \pm 0.16$ | $0.48 \pm 0.03$ | $0.70 \pm 0.20$ | $0.51 \pm 0.03$ | $0.70 \pm 0.29$ | $0.61 \pm 0.24$ | $0.52 \pm 0.16$ | $0.55 \pm 0.19$ | $0.54 \pm 0.19$ | $0.501 \pm 0.007$ |
| Log. regr. | $0.607 \pm 0.058$ | $1.00 \pm 0.00$ | $0.54 \pm 0.04$ | $0.99 \pm 0.00$ | $0.51 \pm 0.04$ | $1.00 \pm 0.00$ | $1.00 \pm 0.00$ | $0.93 \pm 0.15$ | $0.60 \pm 0.19$ | $0.43 \pm 0.16$ | $0.826 \pm 0.211$ |
| Burden test | $0.699 \pm 0.041$ | $1.00 \pm 0.00$ | $0.64 \pm 0.05$ | $1.00 \pm 0.00$ | $0.89 \pm 0.02$ | $1.00 \pm 0.00$ | $1.00 \pm 0.00$ | $1.00 \pm 0.00$ | $1.00 \pm 0.00$ | $0.79 \pm 0.28$ | $0.549 \pm 0.074$ |
| Log. MIL (KMER) | $0.582 \pm 0.065$ | $0.54 \pm 0.07$ | $0.51 \pm 0.03$ | $0.99 \pm 0.00$ | $0.62 \pm 0.15$ | $1.00 \pm 0.00$ | $0.72 \pm 0.11$ | $0.64 \pm 0.14$ | $0.57 \pm 0.15$ | $0.53 \pm 0.13$ | $0.665 \pm 0.224$ |
| Log. MIL (TCRβ) | $0.515 \pm 0.073$ | $0.50 \pm 0.03$ | $0.50 \pm 0.02$ | $0.99 \pm 0.00$ | $0.78 \pm 0.03$ | $0.54 \pm 0.09$ | $0.57 \pm 0.16$ | $0.47 \pm 0.09$ | $0.51 \pm 0.07$ | $0.50 \pm 0.12$ | $0.501 \pm 0.016$ |
| Known motif b. | – | $1.00 \pm 0.00$ | $0.70 \pm 0.03$ | $0.99 \pm 0.00$ | $0.62 \pm 0.04$ | $1.00 \pm 0.00$ | $1.00 \pm 0.00$ | $1.00 \pm 0.00$ | $1.00 \pm 0.00$ | $1.00 \pm 0.00$ | $0.890 \pm 0.168$ |
| Known motif c. | – | $0.92 \pm 0.00$ | $0.56 \pm 0.03$ | $0.65 \pm 0.03$ | $0.52 \pm 0.03$ | $1.00 \pm 0.00$ | $1.00 \pm 0.00$ | $0.99 \pm 0.01$ | $0.72 \pm 0.09$ | $0.63 \pm 0.09$ | $0.738 \pm 0.202$ |

Our proposed method, DeepRC, also uses a MIL approach but considers sequences rather than k-mers as instances within an input object and a transformer-like attention mechanism. DeepRC sets out to avoid the above-mentioned constraints of current methods by (a) applying transformer-like attention-pooling instead of max-pooling and learning a classifier on the repertoire rather than on the sequence-representation, (b) pooling learned representations rather than predictions, and (c) using less rigid feature extractors, such as 1D convolutions or LSTMs. *In this work, we contribute the following:* We demonstrate that continuous generalizations of binary modern Hopfield-networks (Krotov & Hopfield, 2016, 2018; Demircigil et al., 2017) have an update rule that is known as the attention mechanisms in the transformer. We show that these modern Hopfield networks have exponential storage capacity, which allows them to extract patterns among a large set of instances (next Section). Based on this result, we propose DeepRC, a novel deep MIL method based on modern Hopfield networks for large bags of complex sequences, as they occur in immune repertoire classification (Section “Deep Repertoire Classification). We evaluate the predictive performance of DeepRC and other machine learning approaches for the classification of immune repertoires in a large comparative study (Section “Experimental Results”)

### Exponential storage capacity of continuous state modern Hopfield networks with transformer attention as update rule

In this section, we show that modern Hopfield networks have exponential storage capacity, which will later allow us to approach massive multiple-instance learning problems, such as immune repertoire classification. See our companion paper (Ramsauer et al., 2020) for a detailed derivation and analysis of modern Hopfield networks.

We assume patterns ***x***_1_, …, ***x***_*N*_ ∈ ℝ^*d*^ that are stacked as columns to the matrix ***X*** = (***x***_1_, …, ***x***_*N*_) and a query pattern ***ξ*** that also represents the current state. The largest norm of a pattern is *M* = max_*i*_ ‖***x***_*i*_‖. The *separation* Δ_*i*_ of a pattern ***x***_*i*_ is defined as its minimal dot product difference to any of the other patterns: 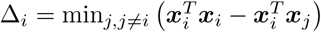. A pattern is *well-separated* from the data if 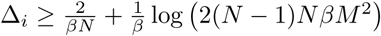. We consider a modern Hopfield network with current state ***ξ*** and the energy function 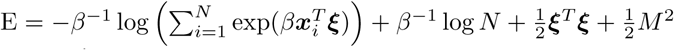. For energy E and state ***ξ***, the update rule

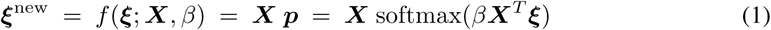

is proven to converge globally to stationary points of the energy E, which are local minima or saddle points (see (Ramsauer et al., 2020), appendix, Theorem A2). *Surprisingly, the update rule Eq*. (1) *is also the formula of the well-known transformer attention mechanism*.

To see this more clearly, we simultaneously update several queries ***ξ***_*i*_. Furthermore the queries ***ξ***_*i*_ and the patterns ***x***_*i*_ are linear mappings of vectors ***y***_*i*_ into the space ℝ^*d*^. For matrix notation, we set 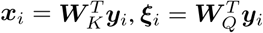 and multiply the result of our update rule with ***W***_*V*_. Using ***Y*** = (***y***_1_, …, ***y***_*N*_)^*T*^, we define the matrices ***X***^*T*^ = ***K*** = ***Y W***_*K*_, ***Q*** = ***Y W***_*Q*_, and ***V*** = ***Y W***_*K*_***W***_*V*_ = ***X***^*T*^ ***W***_*V*_, where 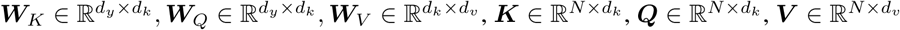, and the patterns are now mapped to the Hopfield space with dimension *d* = *d*_*k*_. We set 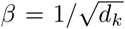 and change softmax to a row vector. The update rule Eq. (1) multiplied by ***W***_*V*_ performed for all queries simultaneously becomes in row vector notation:

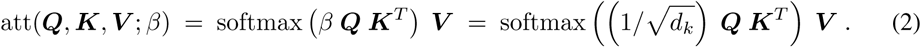

This formula is the transformer attention.

If the patterns ***x***_*i*_ are well separated, the iterate Eq. (1) converges to a fixed point close to a pattern to which the initial ***ξ*** is similar. If the patterns are not well separated the iterate Eq.(1) converges to a fixed point close to the arithmetic mean of the patterns. If some patterns are similar to each other but well separated from all other vectors, then a *metastable state* between the similar patterns exists. Iterates that start near a metastable state converge to this metastable state. For details see Ramsauer et al. (2020), appendix, Sect. A2. Typically, the update converges after one update step (see Ramsauer et al. (2020), appendix, Theorem A8) and has an exponentially small retrieval error (see Ramsauer et al. (2020), appendix, Theorem A9).

Our main concern for application to immune repertoire classification is the number of patterns that can be stored and retrieved by the modern Hopfield network, equivalently to the transformer attention head. The storage capacity of an attention mechanism is critical for massive MIL problems. We first define what we mean by storing and retrieving patterns from the modern Hopfield network.

#### Definition 1

(Pattern Stored and Retrieved). *We assume that around every pattern* ***x***_*i*_ *a sphere* S_*i*_ *is given. We say* ***x***_*i*_ is stored *if there is a single fixed point* 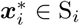 *to which all points* ***ξ*** ∈ S_*i*_ *converge, and* S_*i*_ ∩ S_*j*_ = ∅ *for i* ≠ *j. We say* ***x***_*i*_ is retrieved *if the iteration Eq*. (1) *converged to the single fixed point* 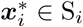.

For randomly chosen patterns, the number of patterns that can be stored is exponential in the dimension *d* of the space of the patterns (***x***_*i*_ ∈ ℝ^*d*^).

#### Theorem 1.

*We assume a failure probability* 0 *< p ⩽* 1 *and randomly chosen patterns on the sphere with radius* 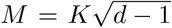. *We define* 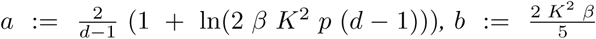, *and* 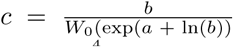, *where W*_0_ *is the upper branch of the Lambert W function and ensure* 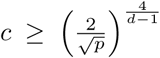. *Then with probability* 1 − *p, the number of random patterns that can be stored is*

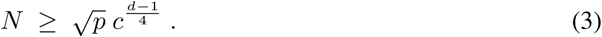

*Examples are c* ≥ 3.1546 *for β* = 1, *K* = 3, *d* = 20 *and p* = 0.001 *(a* + ln(*b*) *>* 1.27*) and c* ≥ 1.3718 *for β* = 1 *K* = 1, *d* = 75, *and p* = 0.001 *(a* + ln(*b*) *<* −0.94).

See Ramsauer et al. (2020), appendix, Theorem A5 for a proof. We have established that a modern Hopfield network or a transformer attention mechanism can store and retrieve exponentially many patterns. This allows us to approach MIL with massive numbers of instances from which we have to retrieve a few with an attention mechanism.

### Deep Repertoire Classification

#### Problem setting and notation

We consider a MIL problem, in which an input object *X* is a *bag* of *N* instances *X* = {*s*_1_, …, *s*_*N*_}. The instances do not have dependencies nor orderings between them and *N* can be different for every object. We assume that each instance *s*_*i*_ is associated with a label *y*_*i*_ ∈ {0, 1}, assuming a binary classification task, to which we do not have access. We only have access to a label *Y* = max_*i*_ *y*_*i*_ for an input object or bag. Note that this poses a credit assignment problem, since the sequences that are responsible for the label *Y* have to be identified and that the relation between instance-label and bag-label can be more complex (Foulds & Frank, 2010).

A model *ŷ* = *g*(*X*) should be (a) invariant to permutations of the instances and (b) able to cope with the fact that *N* varies across input objects (Ilse et al., 2018), which is a problem also posed by point sets (Qi et al., 2017). Two principled approaches exist. The first approach is to learn an instance-level scoring function *h* : 𝒮 ⟼ [0, 1], which is then pooled across instances with a pooling function *f*, for example by average-pooling or max-pooling (see below). The second approach is to construct an instance representation ***z***_*i*_ of each instance by 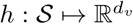 and then encode the bag, or the input object, by pooling these instance representations (Wang et al., 2018) via a function *f*. An output function 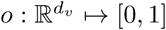 subsequently classifies the bag. The second approach, the pooling of representations rather than scoring functions, is currently best performing (Wang et al., 2018).

In the problem at hand, the input object *X* is the immune repertoire of an individual that consists of a large set of immune receptor sequences (T-cell receptors or antibodies). Immune receptors are primarily represented as sequences *s*_*i*_ from a space *s*_*i*_ ∈ 𝒮. These sequences act as the instances in the MIL problem. Although immune repertoire classification can readily be formulated as a MIL problem, it is yet unclear how well machine learning methods solve the above-described problem with a large number of instances *N* ≫ 10, 000 and with instances *s*_*i*_ being complex sequences. Next we describe currently used pooling functions for MIL problems.

#### Pooling functions for MIL problems

Different pooling functions equip a model *g* with the property to be invariant to permutations of instances and with the ability to process different numbers of instances. Typically, a neural network *h*_***θ***_ with parameters ***θ*** is trained to obtain a function that maps each instance onto a representation: ***z***_*i*_ = *h*_***θ***_(*s*_*i*_) and then a pooling function ***z*** = *f* ({***z***_1_, …, ***z***_*N*_}) supplies a representation ***z*** of the input object *X* = {*s*_1_, …, *s*_*N*_}. The following pooling functions are typically used: *average-pooling*: 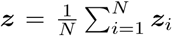, *max-pooling*: 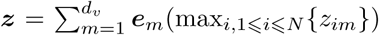, where ***e***_*m*_ is the standard basis vector for dimension *m* and *attention-pooling*: 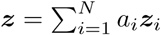, where *a*_*i*_ are non-negative (*a*_*i*_ ≥ 0), sum to one 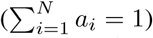, and are determined by an attention mechanism. These pooling functions are invariant to permutations of {1, …, *N*} and are differentiable. Therefore, they are suited as building blocks for Deep Learning architectures. We employ attention-pooling in our DeepRC model as detailed in the following.

#### Modern Hopfield networks viewed as transformer-like attention mechanisms

The modern Hop-field networks, as introduced above,have a storage capacity that is exponential in the dimension of the vector space and converge after just one update (see (Ramsauer et al., 2020), appendix).Additionally, the update rule of modern Hopfield networks is known as key-value attention mechanism, which has been highly successful through the transformer (Vaswani et al., 2017) and BERT (Devlin et al., 2019) models in natural language processing. Therefore using modern Hopfield networks with the key-value-attention mechanism as update rule is the natural choice for our task. In particular, modern Hopfield networks are theoretically justified for storing and retrieving the large number of vectors (sequence patterns) that appear in the immune repertoire classification task.

Instead of using the terminology of modern Hopfield networks, we explain our DeepRC architecture in terms of key-value-attention (the update rule of the modern Hopfield network), since it is well known in the deep learning community. The attention mechanism assumes a space of dimension *d*_*k*_ in which keys and queries are compared. A set of *N* key vectors are combined to the matrix ***K***. A set of *d*_*q*_ query vectors are combined to the matrix ***Q***. Similarities between queries and keys are computed by inner products, therefore queries can search for similar keys that are stored. Another set of *N* value vectors are combined to the matrix ***V***. The output of the attention mechanism is a weighted average of the value vectors for each query ***q***. The *i*-th vector ***v***_*i*_ is weighted by the similarity between the *i*-th key ***k***_*i*_ and the query ***q***. The similarity is given by the softmax of the inner products of the query ***q*** with the keys ***k***_*i*_. All queries are calculated in parallel via matrix operations. Consequently, the attention function att(***Q, K, V***; *β*) maps queries ***Q***, keys ***K***, and values ***V*** to *d*_*v*_-dimensional outputs: att(***Q, K, V***; *β*) = softmax(*β****QK***^*T*^)***V*** (see also Eq. (2)). While this attention mechanism has originally been developed for sequence tasks (Vaswani et al., 2017), it can be readily transferred to sets (Lee et al., 2019; Ye et al., 2018). This type of attention mechanism will be employed in DeepRC.

#### The DeepRC method

We propose a novel method **Deep R**epertoire **C**lassification (**DeepRC**) for immune repertoire classification with attention-based deep massive multiple instance learning and compare it against other machine learning approaches. For DeepRC, we consider *immune repertoires as input objects*, which are represented as bags of instances. In a bag, *each instance is an immune receptor sequence* and each bag can contain a large number of sequences. Note that we will use ***z***_*i*_ to denote the *sequence-representation* of the *i*-th sequence and ***z*** to denote the *repertoire-representation*. At the core, DeepRC consists of a transformer-like attention mechanism that extracts the most important information from each repertoire. We first give an overview of the attention mechanism and then provide details on each of the sub-networks *h*_1_, *h*_2_, and *o* of DeepRC. (Overview: Fig. 1; Architecture: Fig. 2; Implementation details: Sect. A2; DeepRC variations: Sect. A10.)

**Figure 1:**
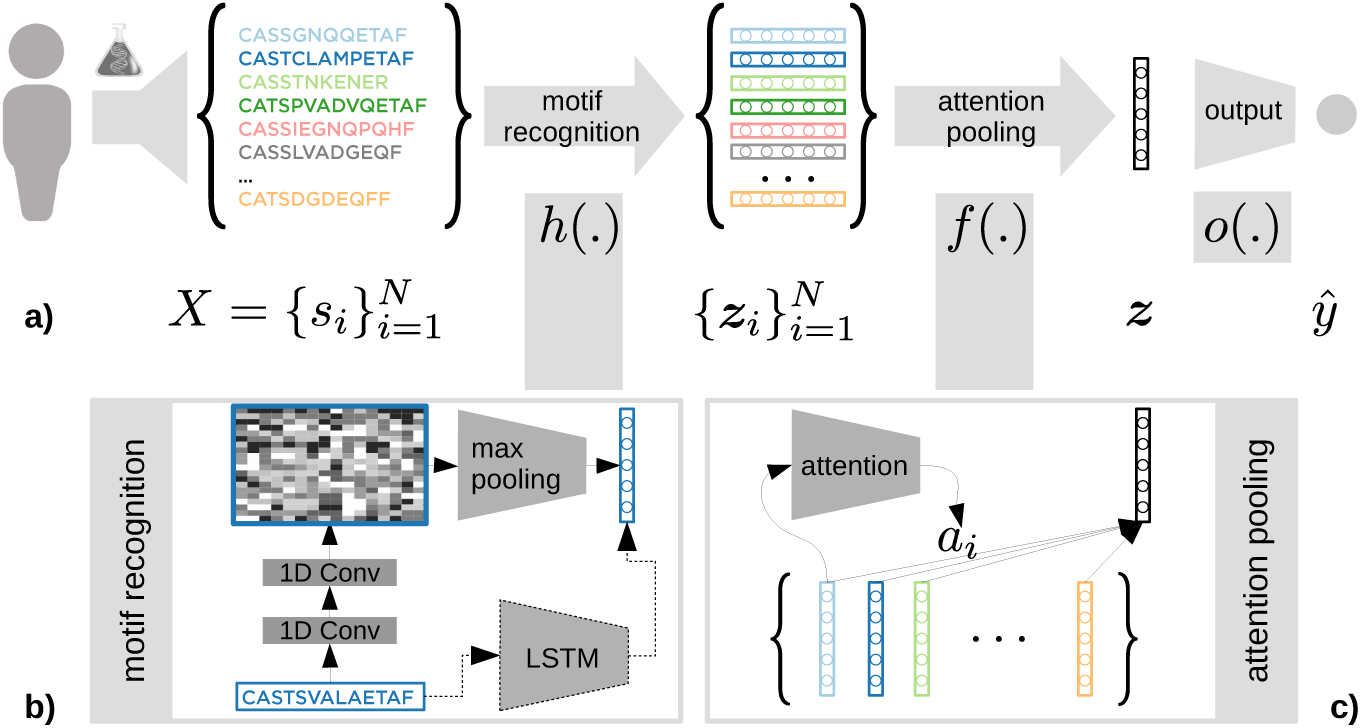
Schematic representation of the DeepRC approach. **a)** An immune repertoire *X* is represented by large bags of immune receptor sequences (colored). A neural network (NN) *h* serves to recognize patterns in each of the sequences *s*_*i*_ and maps them to sequence-representations ***z***_*i*_. A pooling function *f* is used to obtain a repertoire-representation ***z*** for the input object. Finally, an output network *o* predicts the class label *ŷ* **b)** DeepRC uses stacked 1D convolutions for a parameterized function *h* due to their computational efficiency. Potentially, millions of sequences have to be processed for each input object. In principle, also recurrent neural networks (RNNs), such as LSTMs (Hochreiter et al., 2007), or transformer networks (Vaswani et al., 2017) may be used but are currently computationally too costly. **c)** Attention-pooling is used to obtain a repertoire-representation ***z*** for each input object, where DeepRC uses weighted averages of sequence-representations. The weights are determined by an update rule of modern Hopfield networks that allows to retrieve exponentially many patterns.

**Figure 2:**
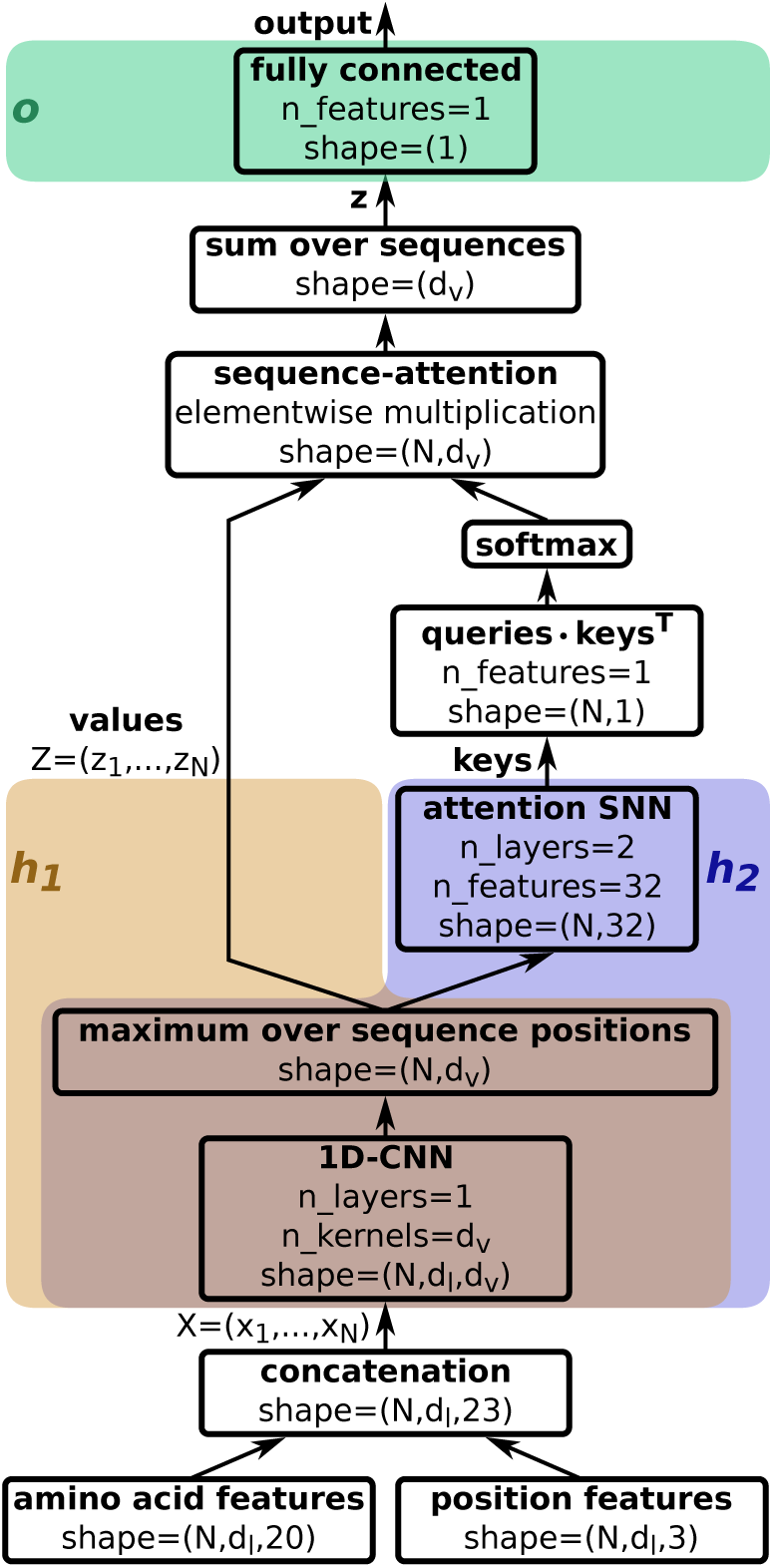
DeepRC architecture as used in Table 1 with sub-networks *h*_1_, *h*_2_, and *o. d*_*l*_ indicates the sequence length.

#### Attention mechanism in DeepRC

This mechanism is based on the three matrices ***K*** (the keys), ***Q*** (the queries), and ***V*** (the values) together with a parameter *β. Values*. DeepRC uses a 1D convolutional network *h*_1_ (LeCun et al., 1998; Hu et al., 2014; Kelley et al., 2016) that supplies a sequence-representation ***z***_*i*_ = *h*_1_(*s*_*i*_), which acts as the values ***V*** = ***Z*** = (***z***_1_, …, ***z***_*N*_) in the attention mechanism (see Figure 2). *Keys*. A second neural network *h*_2_, which shares its first layers with *h*_1_, is used to obtain keys 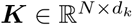 for each sequence in the repertoire. This network uses 2 self-normalizing layers (Klambauer et al., 2017) with 32 units per layer (see Figure 2). *Query*. We use a fixed *d*_*k*_-dimensional query vector ***ξ*** which is learned via backpropagation. For more attention heads, each head has a fixed query vector. With the quantities introduced above, the transformer attention mechanism (Eq. (2)) of DeepRC is implemented as follows:

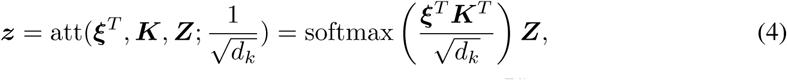

where 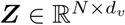 are the sequence–representations stacked row-wise, ***K*** are the keys, and ***z*** is the repertoire-representation and at the same time a weighted mean of sequence–representations ***z***_*i*_. The attention mechanism can readily be extended to multiple queries, however, computational demand could constrain this depending on the application and dataset. Theorem 1 demonstrates that this mechanism is able to retrieve a single pattern out of several hundreds of thousands.

##### Attention-pooling and interpretability

Each input object, i.e. repertoire, consists of a large number *N* of sequences, which are reduced to a single fixed-size feature vector of length *d*_*v*_ representing the whole input object by an attention-pooling function. To this end, a transformer-like attention mechanism adapted to sets is realized in DeepRC which supplies *a*_*i*_ – the importance of the sequence *s*_*i*_. This importance value is an interpretable quantity, which is highly desired for the immunological problem at hand. Thus, DeepRC allows for two forms of interpretability methods. (a) A trained DeepRC model can compute attention weights *a*_*i*_, which directly indicate the importance of a sequence. (b) DeepRC furthermore allows for the usage of contribution analysis methods, such as Integrated Gradients (IG) (Sundararajan et al., 2017) or Layer-Wise Relevance Propagation (Montavon et al., 2018; Arras et al., 2019). See Sect. A8 for details.

#### Classification layer and network parameters

The repertoire-representation ***z*** is then used as input for a fully-connected output network *ŷ* = *o*(***z***) that predicts the immune status, where we found it sufficient to train single-layer networks. In the simplest case, DeepRC predicts a single target, the class label *y*, e.g. the immune status of an immune repertoire, using one output value. However, since DeepRC is an end-to-end deep learning model, multiple targets may be predicted simultaneously in classification or regression settings or a mix of both. This allows for the introduction of additional information into the system via auxiliary targets such as age, sex, or other metadata.

#### Network parameters, training, and inference

DeepRC is trained using standard gradient descent methods to minimize a cross-entropy loss. The network parameters are ***θ***_1_, ***θ***_2_, ***θ***_*o*_ for the sub-networks *h*_1_, *h*_2_, and *o*, respectively, and additionally ***ξ***. In more detail, we train DeepRC using Adam (Kingma & Ba, 2014) with a batch size of 4 and dropout of input sequences.

#### Implementation

To reduce computational time, the attention network first computes the attention weights *a*_*i*_ for each sequence *s*_*i*_ in a repertoire. Subsequently, the top 10% of sequences with the highest *a*_*i*_ per repertoire are used to compute the weight updates and prediction. Furthermore, computation of ***z***_*i*_ is performed in 16-bit, others in 32-bit precision to ensure numerical stability in the softmax. See Sect. A2 for details.

### Experimental Results

In this section, we report and analyze the predictive power of DeepRC and the compared methods on several immunosequencing datasets. The ROC-AUC is used as the main metric for the predictive power.

#### Methods compared

We compared previous methods for immune repertoire classification, (Ostmeyer et al., 2019) (“Log. MIL (KMER)”, “Log. MIL (TCRB)”) and a burden test (Emerson et al., 2017), as well as the baseline methods Logistic Regression (“Log. Regr.”), k-nearest neighbour (“KNN”), and Support Vector Machines (“SVM”) with kernels designed for sets, such as the Jaccard kernel (“J”) and the MinMax (“MM”) kernel (Ralaivola et al., 2005). For the simulated data, we also added baseline methods that search for the implanted motif either in binary or continuous fashion (“Known motif b.”, “Known motif c.”) assuming that this motif was known (for details, see Sect. A4).

#### Datasets

We aimed at constructing immune repertoire classification scenarios with varying degree of difficulties and realism in order to compare and analyze the suggested machine learning methods. To this end, we either use simulated or experimentally-observed immune receptor sequences and we implant signals, specifically, sequence motifs or sets thereof (Akbar et al., 2019; Weber et al., 2020), at different frequencies into sequences of repertoires of the positive class. These frequencies represent the *witness rates* and range from 0.01% to 10%. Overall, we compiled four categories of datasets: (a) simulated immunosequencing data with implanted signals, (b) LSTM-generated immunosequencing data with implanted signals, (c) real-world immunosequencing data with implanted signals, and (d) real-world immunosequencing data with known immune status, the *CMV dataset* (Emerson et al., 2017). The average number of instances per bag, which is the number of sequences per immune repertoire, is ≈300,000 except for category (c), in which we consider the scenario of low-coverage data with only 10,000 sequences per repertoire. The number of repertoires per dataset ranges from 785 to 5,000. In total, all datasets comprise ≈30 billion sequences or instances. This represents the largest comparative study on immune repertoire classification (see Sect. A3).

#### Hyperparameter selection

We used a nested 5-fold cross validation (CV) procedure to estimate the performance of each of the methods. All methods could adjust their most important hyperparameters on a validation set in the inner loop of the procedure. See Sect. A5 for details.

#### Results

In each of the four categories, “real-world data”, “real-world data with implanted signals”, “LSTM-generated data”, and “simulated immunosequencing data”, DeepRC outperforms all competing methods with respect to average AUC. Across categories, the runner-up methods are either the SVM for MIL problems with MinMax kernel or the burden test (see Table 1 and Sect. A6).

##### Results on simulated immunosequencing data

In this setting the complexity of the implanted signal is in focus and varies throughout 18 simulated datasets (see Sect. A3). Some datasets are challenging for the methods because the implanted motif is hidden by noise and others because only a small fraction of sequences carries the motif, and hence have a low witness rate. These difficulties become evident by the method called “known motif binary”, which assumes the implanted motif is known. The performance of this method ranges from a perfect AUC of 1.000 in several datasets to an AUC of 0.532 in dataset ‘17’ (see Sect. A6). DeepRC outperforms all other methods with an average AUC of 0.846 ± 0.223, followed by the SVM with MinMax kernel with an average AUC of 0.827 ± 0.210 (see Sect. A6). The predictive performance of all methods suffers if the signal occurs only in an extremely small fraction of sequences. In datasets, in which only 0.01% of the sequences carry the motif, all AUC values are below 0.550. *Results on LSTM-generated data*. On the LSTM-generated data, in which we implanted noisy motifs with frequencies of 10%, 1%, 0.5%, 0.1%, and 0.05%, DeepRC yields almost perfect predictive performance with an average AUC of 1.000 ± 0.001 (see Sect. A6 and A7). The second best method, SVM with MinMax kernel, has a similar predictive performance to DeepRC on all datasets but the other competing methods have a lower predictive performance on datasets with low frequency of the signal (0.05%). *Results on real-world data with implanted motifs*. In this dataset category, we used real immunosequences and implanted single or multiple noisy motifs. Again, DeepRC outperforms all other methods with an average AUC of 0.980 ± 0.029, with the second best method being the burden test with an average AUC of 0.883 ± 0.170. Notably, all methods except for DeepRC have difficulties with noisy motifs at a frequency of 0.1% (see Tab. A11). *Results on real-world data*. On the real-world dataset, in which the immune status of persons affected by the cytomegalovirus has to be predicted, the competing methods yield predictive AUCs between 0.515 and 0.825 (see Table 1). We note that this dataset is not the exact dataset that was used in Emerson et al. (2017). It differs in pre-processing and also comprises a different set of samples and a smaller training set due to the nested 5-fold cross-validation procedure, which leads to a more challenging dataset. The best performing method is DeepRC with an AUC of 0.831 ± 0.002, followed by the SVM with MinMax kernel (AUC 0.825 ± 0.022) and the burden test with an AUC of 0.699 ± 0.041. The top-ranked sequences by DeepRC significantly correspond to those detected by Emerson et al. (2017), which we tested by a Mann-Whitney U-test with the null hypothesis that the attention values of the sequences detected by Emerson et al. (2017) would be equal to the attention values of the remaining sequences (*p*-value of 1.3 · 10^−93^). The sequence attention values are displayed in Tab. A14.

#### Conclusion

We have demonstrated how modern Hopfield networks and attention mechanisms enable successful classification of the immune status of immune repertoires. For this task, methods have to identify the discriminating sequences amongst a large set of sequences in an immune repertoire. Specifically, even motifs within those sequences have to be identified. We have shown that DeepRC, a modern Hopfield network and an attention mechanism with a fixed query, can solve this difficult task despite the massive number of instances. DeepRC furthermore outperforms the compared methods across a range of different experimental conditions.

### Broader Impact

#### Impact on machine learning and related scientific fields

We envision that with (a) the increasing availability of large immunosequencing datasets (Kovaltsuk et al., 2018; Corrie et al., 2018; Christley et al., 2018; Zhang et al., 2020; Rosenfeld et al., 2018; Shugay et al., 2018), (b) further fine-tuning of ground-truth benchmarking immune receptor datasets (Weber et al., 2020; Olson et al., 2019; Marcou et al., 2018), (c) accounting for repertoire-impacting factors such as age, sex, ethnicity, and environment (potential confounding factors), and (d) increased GPU memory and increased computing power, it will be possible to identify discriminating immune receptor motifs for many diseases, potentially even for the current SARS-CoV-2 (COVID-19) pandemic (Raybould et al., 2020; Minervina et al., 2020; Galson et al., 2020). Such results would greatly benefit ongoing research on antibody and TCR-driven immunotherapies and immunodiagnostics as well as rational vaccine design (Brown et al., 2019).

In the course of this development, the experimental verification and interpretation of machine-learning-identified motifs could receive additional focus, as for most of the sequences within a repertoire the corresponding antigen is unknown. Nevertheless, recent technological breakthroughs in high-throughput antigen-labeled immunosequencing are beginning to generate large-scale antigen-labeled single-immune-receptor-sequence data thus resolving this longstanding problem (Setliff et al., 2019).

From a machine learning perspective, the successful application of DeepRC on immune repertoires with their large number of instances per bag might encourage the application of modern Hopfield networks and attention mechanisms on new, previously unsolved or unconsidered, datasets and problems.

#### Impact on society

If the approach proves itself successful, it could lead to faster testing of individuals for their immune status w.r.t. a range of diseases based on blood samples. This might motivate changes in the pipeline of diagnostics and tracking of diseases, e.g. automated testing of the immune status in regular intervals. It would furthermore make the collection and screening of blood samples for larger databases more attractive. In consequence, the improved testing of immune statuses might identify individuals that do not have a working immune response towards certain diseases to government or insurance companies, which could then push for targeted immunisation of the individual. Similarly to compulsory vaccination, such testing for the immune status could be made compulsory by governments, possibly violating privacy or personal self-determination in exchange for increased over-all health of a population.

Ultimately, if the approach proves itself successful, the insights gained from the screening of individuals that have successfully developed resistances against specific diseases could lead to faster targeted immunisation, once a certain number of individuals with resistances can be found. This might strongly decrease the harm done by e.g. pandemics and lead to a change in the societal perception of such diseases.

#### Consequences of failures of the method

As common with methods in machine learning, potential danger lies in the possibility that users rely too much on our new approach and use it without reflecting on the outcomes. However, the full pipeline in which our method would be used includes wet lab tests after its application, to verify and investigate the results, gain insights, and possibly derive treatments. Failures of the proposed method would lead to unsuccessful wet lab validation and negative wet lab tests. Since the proposed algorithm does not directly suggest treatment or therapy, human beings are not directly at risk of being treated with a harmful therapy. Substantial wet lab and in-vitro testing and would indicate wrong decisions by the system.

#### Leveraging of biases in the data and potential discrimination

As for almost all machine learning methods, confounding factors, such as age or sex, could be used for classification. This, might lead to biases in predictions or uneven predictive performance across subgroups. As a result, failures in the wet lab would occur (see paragraph above). Moreover, insights into the relevance of the confounding factors could be gained, leading to possible therapies or counter-measures concerning said factors.

Furthermore, the amount of data available with respec to relevant confounding factors could lead to better or worse performance of our method. E.g. a dataset consisting mostly of data from individuals within a specific age group might yield better performance for that age group, possibly resulting in better or exclusive treatment methods for that specific group. Here again, the application of DeepRC would be followed by in-vitro testing and development of a treatment, where all target groups for the treatment have to be considered accordingly.

## Availability

All datasets and code is available at https://github.com/ml-jku/DeepRC. The CMV dataset is publicly available at https://clients.adaptivebiotech.com/pub/Emerson-2017-NatGen.

## Acknowledgments

The ELLIS Unit Linz, the LIT AI Lab and the Institute for Machine Learning are supported by the Land Oberösterreich, LIT grants DeepToxGen (LIT-2017-3-YOU-003), and AI-SNN (LIT-2018-6-YOU-214), the Medical Cognitive Computing Center (MC3), Janssen Pharmaceutica, UCB Biopharma, Merck Group, Audi.JKU Deep Learning Center, Audi Electronic Venture GmbH, TGW, Primal, Silicon Austria Labs (SAL), FILL, EnliteAI, Google Brain, ZF Friedrichshafen AG, Robert Bosch GmbH, TÜV Austria, DCS, and the NVIDIA Corporation. Victor Greiff (VG) and Geir Kjetil Sandve (GKS) are supported by The Helmsley Charitable Trust (#2019PG-T1D011, to VG), UiO World-Leading Research Community (to VG), UiO:LifeSciences Convergence Environment Immunolingo (to VG and GKS), EU Horizon 2020 iReceptorplus (#825821, to VG) and Stiftelsen Kristian Gerhard Jebsen (K.G. Jebsen Coeliac Disease Research Centre, to GKS).

## Appendix

In the following, the appendix to the paper “Modern Hopfield Networks and Attention for Immune Repertoire Classification” is presented. Here we provide details on DeepRC, the compared methods, and the experimental setup and results. Furthermore, the generation of the immune repertoire classification data using an LSTM network, the interpretation of DeepRC and the extraction of found motifs, and the ablation study using different variants of DeepRC are described.

### A1 Introduction

In Section A2 we provide details on the architecture of DeepRC, in Section A3 we present details on the datasets, in Section A4 we explain the methods that we compared, in Section A5 we elaborate on the hyperparameters and their selection process. Then, in Section A6 we present detailed results for each dataset category in tabular form, in Section A7 we provide information on the LSTM model that was used to generate antibody sequences, in Section A8 we show how DeepRC can be interpreted, in Section A9 we show the correspondence of previously identified TCR sequences for CMV immune status with attention values by DeepRC, and finally we present variations and an ablation study of DeepRC in Section A10.

### A2 DeepRC implementation details

#### Input layer

For the input layer of the CNN, the characters in the input sequence, i.e. the amino acids (AAs), are encoded in a one-hot vector of length 20. To also provide information about the position of an AA in the sequence, we add 3 additional input features with values in range [0, 1] to encode the position of an AA relative to the sequence. These 3 positional features encode whether the AA is located at the beginning, the center, or the end of the sequence, respectively, as shown in Figure A1. We concatenate these 3 positional features with the one-hot vector of AAs, which results in a feature vector of size 23 per sequence position. Each repertoire, now represented as a bag of feature vectors, is then normalized to unit variance. Since the cytomegalovirus dataset (*CMV dataset*) provides sequences with an associated abundance value per sequence, which is the number of occurrences of a sequence in a repertoire, we incorporate this information into the input of DeepRC. To this end, the one-hot AA features of a sequence are multiplied by a scaling factor of log(*c*_*a*_) before normalization, where *c*_*a*_ is the abundance of a sequence. We feed the sequences with 23 features per position into the CNN. Sequences of different lengths were zero-padded to the maximum sequence length per batch at the sequence ends.

#### 1D CNN for motif recognition

In the following, we describe how DeepRC identifies patterns in the individual sequences and reduces each sequence in the input object to a fixed-size feature vector. DeepRC employs 1D convolution layers to extract patterns, where trainable weight kernels are convolved over the sequence positions. In principle, also recurrent neural networks (RNNs) or transformer networks could be used instead of 1D CNNs, however, (a) the computational complexity of the network must be low to be able to process millions of sequences for a single update. Additionally, (b) the learned network should be able to provide insights in the recognized patterns in form of motifs. Both properties (a) and (b) are fulfilled by 1D convolution operations that are used by DeepRC.

We use one 1D CNN layer (Hu et al., 2014) with SELU activation functions (Klambauer et al., 2017) to identify the relevant patterns in the input sequences with a computationally light-weight operation. The larger the kernel size, the more surrounding sequence positions are taken into account, which influences the length of the motifs that can be extracted. We therefore adjust the kernel size during hyperparameter search. In prior works (Ostmeyer et al., 2019), a k-mer size of 4 yielded good predictive performance, which could indicate that a kernel size in the range of 4 may be a proficient choice. For *d*_*v*_ trainable kernels, this produces a feature vector of length *d*_*v*_ at each sequence position. Subsequently, global max-pooling over all sequence positions of a sequence reduces the sequence-representations ***z***_*i*_ to vectors of the fixed length *d*_*v*_. Given the challenging size of the input data per repertoire, the computation of the CNN activations and weight updates is performed using 16-bit floating point values. A list of hyperparameters evaluated for DeepRC is given in Table A3. A comparison of RNN-based and CNN-based sequence embedding for motif recognition in a smaller experimental setting is given in Sec. A10.

**Figure A1:**
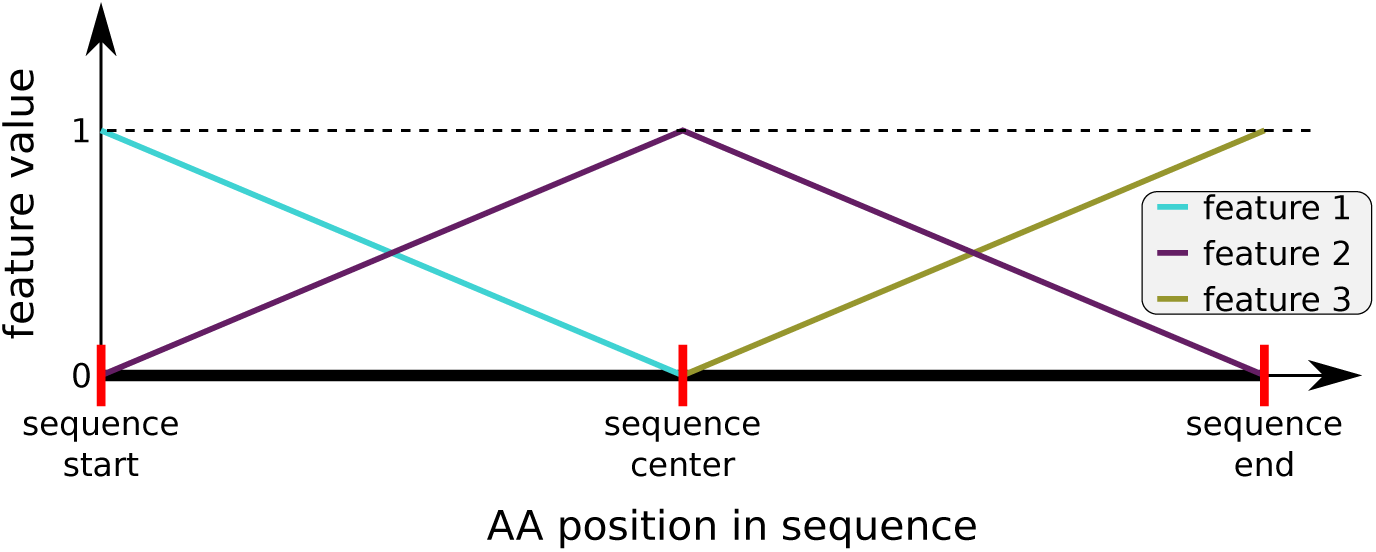
We use 3 input features with values in range [0, 1] to encode the relative position of each AA in a sequence with respect to the sequence. “feature 1” encodes if an AA is close to the sequence start, “feature 2” to the sequence center, and “feature 3” to the sequence end. For every position in the sequence, the values of all three features sum up to 1.

#### Regularization

We apply random and attention-based subsampling of repertoire sequences to reduce over-fitting and decrease computational effort. During training, each repertoire is subsampled to 10, 000 input sequences, which are randomly drawn from the respective repertoire. This can also be interpreted as random drop-out (Hinton et al., 2012) on the input sequences or attention weights. During training and evaluation, the attention weights computed by the attention network are furthermore used to rank the input sequences. Based on this ranking, the repertoire is reduced to the 10% of sequences with the highest attention weights. These top 10% of sequences are then used to compute the weight updates and the prediction for the repertoire. Additionally, one might employ further regularization techniques, which we only partly investigated further in a smaller experimental setting in Sec. A10 due to high computational demands. Such regularization techniques include *l*1 and *l*2 weight decay, noise in the form of random AA permutations in the input sequences, noise on the attention weights, or random shuffling of sequences between repertoires that belong to the negative class. The last regularization technique assumes that the sequences in positive-class repertoires carry a signal, such as an AA motif corresponding to an immune response, whereas the sequences in negative-class repertoires do not. Hence, the sequences can be shuffled randomly between negative class repertoires without obscuring the signal in the positive class repertoires.

#### Hyperparameters

For the hyperparameter search of DeepRC for the category “simulated immunosequencing data”, we only conducted a full hyperparameter search on the more difficult datasets with motif implantation probabilities below 1%, as described in Table A3. This process was repeated for all 5 folds of the 5-fold cross-validation (CV) and the average score on the 5 test sets constitutes the final score of a method.

Table A3 provides an overview of the hyperparameter search, which was conducted as a grid search for each of the datasets in a nested 5-fold CV procedure, as described in Section A4.

#### Computation time and optimization

We took measures on the implementation level to address the high computational demands, especially GPU memory consumption, in order to make the large number of experiments feasible. We train the DeepRC model with a small batch size of 4 samples and perform computation of inference and updates of the 1D CNN using 16-bit floating point values. The rest of the network is trained using 32-bit floating point values. The Adam parameter for numerical stability was therefore increased from the default value of *ϵ* = 10^−8^ to *ϵ* = 10^−4^. Training was performed on various GPU types, mainly `NVIDIA RTX 2080 Ti` Computation times were highly dependent on the number of sequences in the repertoires and the number and sizes of CNN kernels. A single update on an `NVIDIA RTX 2080 Ti` GPU took approximately 0.0129 to 0.0135 seconds, while requiring approximately 8 to 11 GB GPU memory. Taking these optimizations and GPUs with larger memory (≥ 16 GB) into account, it is already possible to train DeepRC, possibly with multi-head attention and a larger network architecture, on larger datasets (see Sec. A10). Our network implementation is based on `PyTorch 1.3.1` (Paszke et al., 2019).

#### Incorporation of additional inputs and metadata

Additional metadata in the form of sequence-level or repertoire-level features could be incorporated into the input via concatenation with the feature vectors that result from taking the maximum of the 1D CNN outputs w.r.t. the sequence positions. This has the benefit that the attention mechanism and output network can utilize the sequence-level or repertoire-level features for their predictions. Sparse metadata or metadata that is only available during training could be used as auxiliary targets to incorporate the information via gradients into the DeepRC model.

#### Limitations

The current methods are mostly limited by computational complexity, since both hyperparameter and model selection is computationally demanding. For hyperparameter selection, a large number of hyperparameter settings have to be evaluated. For model selection, a single repertoire requires the propagation of many thousands of sequences through a neural network and keeping those quantities in GPU memory in order to perform the attention mechanism and weight update. Thus, increased GPU memory would significantly boost our approach. Increased computational power would also allow for more advanced architectures and attention mechanisms, which may further improve predictive performance. Another limiting factor is over-fitting of the model due to the currently relatively small number of samples (bags) in real-world immunosequencing datasets in comparison to the large number of instances per bag and features per instance.

### A3 Datasets

We aimed at constructing immune repertoire classification scenarios with varying degree of realism and difficulties in order to compare and analyze the suggested machine learning methods. To this end, we either use simulated or experimentally-observed immune receptor sequences and we implant signals, which are sequence motifs (Akbar et al., 2019; Weber et al., 2020), into sequences of repertoires of the positive class. It has been shown previously that interaction of immune receptors with antigens occur via short sequence stretches (Akbar et al., 2019). Thus, implantation of short motif sequences simulating an immune signal is biologically meaningful. Our benchmarking study comprises four different categories of datasets: (a) Simulated immunosequencing data with implanted signals (where the signal is defined as sets of motifs), (b) LSTM-generated immunosequencing data with implanted signals, (c) real-world immunosequencing data with implanted signals, and (d) real-world immunosequencing data. Each of the first three categories consists of multiple datasets with varying difficulty depending on the type of the implanted signal and the ratio of sequences with the implanted signal. The ratio of sequences with the implanted signal, where each sequence carries at most 1 implanted signal, corresponds to the *witness rate* (WR). We consider binary classification tasks to simulate the immune status of healthy and diseased individuals. We randomly generate immune repertoires with varying numbers of sequences, where we implant sequence motifs in the repertoires of the diseased individuals, i.e. the positive class. The sequences of a repertoire are also randomly generated by different procedures (detailed below). Each sequence is composed of 20 different characters, corresponding to amino acids, and has an average length of 14.5 AAs.

#### A3.1 Simulated immunosequencing data

In the first category, we aim at investigating the impact of the signal frequency, i.e. the WR, and the signal complexity on the performance of the different methods. To this end, we created 18 datasets, whereas each dataset contains a large number of repertoires with a large number of random AA sequences per repertoire. We then implanted signals in the AA sequences of the positive class repertoires, where the 18 datasets differ in frequency and complexity of the implanted signals. In detail, the AAs were sampled randomly independent of their respective position in the sequence, while the frequencies of AAs, distribution of sequence lengths, and distribution of the number of sequences per repertoire, i.e. the number of instances per bag, are following the respective distributions observed in the real-world *CMV dataset* (Emerson et al., 2017). For this, we first sampled the number of sequences for a repertoire from a Gaussian 𝒩 (*µ* = 316*k, σ* = 132*k*) distribution and rounded to the nearest positive integer. We re-sampled if the size was below 5*k*. We then generated random sequences of AAs with a length of 𝒩 (*µ* = 14.5, *σ* = 1.8), again rounded to the nearest positive integers. Each simulated repertoire was then randomly assigned to either the positive or negative class, with 2, 500 repertoires per class. In the repertoires assigned to the positive class, we implanted motifs with an average length of 4 AAs, following the results of the experimental analysis of antigen-binding motifs in antibodies and T-cell receptor sequences by (Akbar et al., 2019). We varied the characteristics of the implanted motifs for each of the 18 datasets with respect to the following parameters: (a) *ρ*, the probability of a motif being implanted in a sequence of a positive repertoire, i.e. the average ratio of sequences containing the motif, which is the witness rate. (b) The number of wildcard positions in the motif. A wildcard position contains a random AA, which is randomly sampled for each sequence. Wildcard positions are located in the center of the implanted motif. (c) The number of deletion positions in the implanted motif. A deletion position has a probability of 0.5 of being removed from the motif. Deletion positions are located in the center of the implanted motifs.

In this way, we generated 18 different datasets of variable difficulty containing in total roughly 28.7 billion sequences. See Table A1 for an overview of the properties of the implanted motifs in the 18 datasets.

#### A3.2 LSTM-generated data

In the second dataset category, we investigate the impact of the signal frequency and complexity in combination with more plausible immune receptor sequences by taking into account the positional AA distributions and other sequence properties. To this end, we trained an LSTM (Hochreiter & Schmidhuber, 1997) in a standard next character prediction (Graves, 2013) setting to create AA sequences with properties similar to experimentally observed immune receptor sequences.

In the first step, the LSTM model was trained on all immuno-sequences in the *CMV dataset* (Emerson et al., 2017) that contain valid information about sequence abundance and have a known CMV label. Such an LSTM model is able to capture various properties of the sequences, including position-dependent probability distributions and combinations, relationships, and order of AAs. We then used the trained LSTM model to generate 1, 000 repertoires in an autoregressive fashion, starting with a start sequence that was randomly sampled from the trained-on dataset. Based on a visual inspection of the frequencies of 4-mers (see Section A7), the similarity of LSTM generated sequences and real sequences was deemed sufficient for the purpose of generating the AA sequences for the datasets in this category. Further details on LSTM training and repertoire generation are given in Section A7.

After generation, each repertoire was assigned to either the positive or negative class, with 500 repertoires per class. We implanted motifs of length 4 with varying properties in the center of the sequences of the positive class to obtain 5 different datasets. Each sequence in the positive repertoires has a probability *ρ* to carry the motif, which was varied throughout 5 datasets and corresponds to the WR (see Table A1). Each position in the motif has a probability of 0.9 to be implanted and consequently a probability of 0.1 that the original AA in the sequence remains, which can be seen as noise on the motif.

#### A3.3 Real-world data with implanted signals

In the third category, we implanted signals into experimentally obtained immuno-sequences, where we considered 4 dataset variations. Each dataset consists of 750 repertoires for each of the two classes, where each repertoire consists of 10*k* sequences. In this way, we aim to simulate datasets with a *low sequencing coverage*, which means that only relatively few sequences per repertoire are available. The sequences were randomly sampled from healthy (CMV negative) individuals from the *CMV dataset* (see below paragraph for explanation). Two signal types were considered: (a) **One signal with one motif**. The AA motif LDR was implanted in a certain fraction of sequences. The pattern is randomly altered at one of the three positions with probabilities 0.2, 0.6, and 0.2, respectively. (b) **One signal with multiple motifs**. One of the three possible motifs `LDR, CAS`, and `GL-N` was implanted with equal probability. Again, the motifs were randomly altered before implantation. The AA motif LDR changed as described above. The AA motif `CAS` was altered at the second position with probability 0.6 and with probability 0.3 at the first position. The pattern `GL-N`, where - denotes a gap location, is randomly altered at the first position with probability 0.6 and the gap has a length of 0, 1, or 2 AAs with equal probability.

**Table A1:** Properties of simulated repertoires, variations of motifs, and motif frequencies, i.e. the witness rate, for the datasets in categories “simulated immunosequencing data”, “LSTM-generated data”, and “real-world data with implanted signals”. Noise types for * are explained in paragraph “real-world data with implanted signals”.

|  | Simulated | LSTM gen. | Real-world |
| --- | --- | --- | --- |
| seq. per bag | $N(316k, 132k)$ | $N(285k, 156k)$ | 10k |
| repertoires | 5,000 | 1,000 | 1,500 |
| motif noise | 0% | 10% | * |
| wildcards | {0; 1; 2} | 0 | 0 |
| deletions | {0; 1} | 0 | 0 |
| mot. freq. $\rho$<br>(in %) | {1; 0.1;<br>0.01} | {10; 1; 0.5;<br>0.1; 0.05} | {1; 0.1} |

Additionally, the datasets differ in the values for *ρ*, the average ratio of sequences carrying a signal, which were chosen as 1% or 0.1%. The motifs were implanted at positions 107, 109, and 114 according to the IMGT numbering scheme for immune receptor sequences (Lefranc et al., 2003) with probabilities 0.3, 0.35 and 0.2, respectively. With the remaining 0.15 chance, the motif is implanted at any other sequence position. This means that the motif occurrence in the simulated sequences is biased towards the middle of the sequence.

#### A3.4 Real-world data: CMV dataset

We used a real-world dataset of 785 repertoires, each of which containing between 4, 371 to 973, 081 (avg. 299, 319) TCR sequences with a length of 1 to 27 (avg. 14.5) AAs, originally collected and provided by Emerson et al. (2017). 340 out of 785 repertoires were labelled as positive for cytomegalovirus (CMV) serostatus, which we consider as the positive class, 420 repertoires with negative CMV serostatus, considered as negative class, and 25 repertoires with unknown status. We changed the number of sequence counts per repertoire from 1 to 1 for 3 sequences. Furthermore, we exclude a total of 99 repertoires with unknown CMV status or unknown information about the sequence abundance within a repertoire, reducing the dataset for our analysis to 686 repertoires, 312 of which with positive and 374 with negative CMV status.

#### A3.5 Comparison to other MIL datasets

We give a non-exhaustive overview of previously considered MIL datasets and problems in Table A2. To our knowledge the datasets considered in this work pose the most challenging MIL problems with respect to the number of instances per bag (column 5).

**Table A2:** MIL datasets with their numbers of bags and numbers of instances. “total number of instances” refers to the total number of instances in the dataset. The simulated and real-world immunosequencing datasets considered in this work contain a by orders of magnitudes larger number of instances per bag than MIL datasets that were considered by machine learning methods up to now.

| Dataset | Total number of bags | Total number of instances | Approx. number of features per instance | Avg. number of instances per bag | Source | Dataset reference |
| --- | --- | --- | --- | --- | --- | --- |
| Simulated immuno-sequencing data (ours) | 5,000 | 1,597,024,310 x 18 datasets | 14.5x20 AA sequence | 316,000 | this work |  |
| LSTM-generated data (ours) | 1,000 | 304,825,671 x 5 datasets | 14.5x20 AA sequence | 285,000 | this work |  |
| Real-world data with implanted signals (ours) | 1,500 | 14,715,421 x 4 datasets | 14.5x20 AA sequence | 10,000 | this work |  |
| CMV (pre-processed by us) | 785 | 234,965,729 | 14.5x20 AA sequence | 299,000 | this work | Emerson et al. (2017) |
| MNIST bags | 50–500 | 500–50,000 | 28x28x1 image | 100 | Ilse et al. (2018) |  |
| Breast Cancer | 58 | approx. 39,000 | 32x32x3 H&E image | 672 | Ilse et al. (2018) | Gelasca et al. (2008) |
| Basal cell carcinomas | 820 | 7,588,767 | 1024x1024x3 H&E image | 9,056 | Kimeswenger et al. (2019) |  |
| Birds | 548 | 10,232 | 38 | 9 | Ruiz et al. (2018) | Briggs et al. (2012) |
| Scene | 2,000 | 18,000 | 15 | 9 | Ruiz et al. (2018) | Zhang & Zhang (2007) |
| Reuters | 2,000 | 7,119 | 243 | 4 | Ruiz et al. (2018) | Sebastiani (2002) |
| CK+ | 430 | 7,915 | 4,391 | 18 | Ruiz et al. (2018) | Lucey et al. (2010) |
| UniProt (Geobacter sulfurreducens) | 379 | 1,250 | 216 | 3 | Ruiz et al. (2018) | Wu et al. (2014) |
| MODIS (aerosol data) | 1,364 | 136,400 | 12 | 100 | Uriot (2019) | https://aeronet.gsfc.nasa.gov |
| MISR1 (aerosol data) | 800 | 80,000 | 16 | 100 | Uriot (2019) | https://aeronet.gsfc.nasa.gov |
| MISR2 (aerosol data) | 800 | 80,000 | 12 | 54 | Uriot (2019) | https://aeronet.gsfc.nasa.gov |
| CORN (crop yield) | 525 | 52,500 | 92 | 100 | Uriot (2019) | https://aeronet.gsfc.nasa.gov |
| WHEAT (crop yield) | 525 | 52,500 | 92 | 100 | Uriot (2019) | https://aeronet.gsfc.nasa.gov |

### A4 Compared methods

We evaluate and compare the performance of DeepRC against a set of machine learning methods that serve as baseline, were suggested, or can readily be adapted to immune repertoire classification. In this section, we describe these compared methods.

#### A4.1 Known motif

This method serves as an estimate for the achievable classification performance using prior knowledge about which motif was implanted. Note that this does not necessarily lead to perfect predictive performance since motifs are implanted with a certain amount of noise and could also be present in the negative class by chance. The *known motif* method counts how often the known implanted motif occurs per sequence for each repertoire and uses this count to rank the repertoires. From this ranking, the Area Under the receiver operator Curve (AUC) is computed as performance measure. Probabilistic AA changes in the known motif are not considered for this count, with the exception of gap positions. We consider two versions of this method: (a) **Known motif binary:** counts the occurrence of the known motif in a sequence and (b) **Known motif continuous:** counts the maximum number of overlapping AAs between the known motif and all sequence positions, which corresponds to a convolution operation with a binary kernel followed by max-pooling. Since the implanted signal is not known in the experimentally obtained *CMV dataset*, this method cannot be applied to this dataset.

#### A4.2 Support Vector Machine (SVM)

The Support Vector Machine (SVM) approach uses a fixed mapping from a bag of sequences to the corresponding k-mer counts. The function *h*_kmer_ maps each sequence *s*_*i*_ to a vector representing the occurrence of k-mers in the sequence. To avoid confusion with the sequence-representation obtained from the CNN layers of DeepRC, we denote ***u***_*i*_ = *h*_kmer_(*s*_*i*_), which is analogous to ***z***_*i*_. Specifically, *u*_*im*_ = (*h*_kmer_(*s*_*i*_))_*m*_ = #{*p*_*m*_ ∈ *s*_*i*_}, where #{*p*_*m*_ ∈ *s*_*i*_} denotes how often the k-mer pattern *p*_*m*_ occurs in sequence *s*_*i*_. Afterwards, average-pooling is applied to obtain 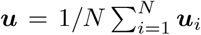, the *k-mer representation* of the input object *X*. For two input objects *X*^(*n*)^ and *X*^(*l*)^ with representations ***u***^(*n*)^ and ***u***^(*l*)^, respectively, we implement the *MinMax kernel* (Ralaivola et al., 2005) as follows:

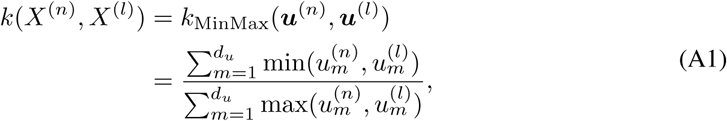

where 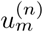 is the *m*-th element of the vector *u*^(*n*)^. The *Jaccard kernel* (Levandowsky & Winter, 1971) is identical to the MinMax kernel except that it operates on binary ***u***^(*n*)^. We used a standard C-SVM, as introduced by Cortes & Vapnik (1995). The corresponding hyperparameter *C* is optimized by random search. The settings of the full hyperparameter search as well as the respective value ranges are given in Table A4a.

#### A4.3 K-Nearest Neighbor (KNN)

The same *k-mer representation* of a repertoire, as introduced above for the SVM baseline, is used for the K-Nearest Neighbor (KNN) approach. As this method clusters samples according to distances between them, the previous kernel definitions cannot be applied directly. It is therefore necessary to transform the MinMax as well as the Jaccard kernel from similarities to distances by constructing the following (Levandowsky & Winter, 1971):

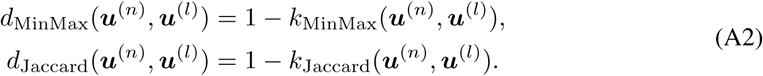

The amount of neighbors is treated as the hyperparameter and optimized by an exhaustive grid search. The settings of the full hyperparameter search as well as the respective value ranges are given in Table A5.

#### A4.4 Logistic regression

We implemented logistic regression on the *k-mer representation* ***u*** of an immune repertoire. The model is trained by gradient descent using the Adam optimizer (Kingma & Ba, 2014). The learning rate is treated as the hyperparameter and optimized by grid search. Furthermore, we explored two regularization settings using combinations of *l*1 and *l*2 weight decay. The settings of the full hyperparameter search as well as the respective value ranges are given in Table A6.

#### A4.5 Burden test

We implemented a burden test (Emerson et al., 2017; Li & Leal, 2008; Wu et al., 2011) in a machine learning setting. The burden test first identifies sequences or k-mers that are associated with the individual’s class, i.e., immune status, and then calculates a burden score per individual. Concretely, for each k-mer or sequence, the phi coefficient of the contingency table for absence or presence and positive or negative immune status is calculated. Then, *J* k-mers or sequences with the highest phi coefficients are selected as the set of associated k-mers or sequences. *J* is a hyperparameter that is selected on a validation set. Additionally, we consider the type of input features, sequences or k-mers, as a hyperparameter. For inference, a burden score per individual is calculated as the sum of associated k-mers or sequences it carries. This score is used as raw prediction and to rank the individuals. Hence, we have extended the burden test by Emerson et al. (2017) to k-mers and to adaptive thresholds that are adjusted on a validation set.

#### A4.6 Logistic MIL (Ostmeyer et al)

The logistic multiple instance learning (MIL) approach for immune repertoire classification (Ostmeyer et al., 2019) applies a logistic regression model to each k-mer representation in a bag. The resulting scores are then summarized by max-pooling to obtain a prediction for the bag. Each amino acid of each k-mer is represented by 5 features, the so-called Atchley factors (Atchley et al., 2005). As k-mers of length 4 are used, this gives a total of 4 ×5 = 20 features. One additional feature per 4-mer is added, which represents the relative frequency of this 4-mer with respect to its containing bag, resulting in 21 features per 4-mer. Two options for the relative frequency feature exist, which are (a) whether the frequency of the 4-mer (“4MER”) or (b) the frequency of the sequence in which the 4-mer appeared (“TCRβ”) is used. We optimized the learning rate, batch size, and early stopping parameter on the validation set. The settings of the full hyperparameter search as well as the respective value ranges are given in Table A8.

### A5 Hyperparameter selection

For all competing methods a hyperparameter search was performed, for which we split each of the 5 training sets into an inner training set and inner validation set. The models were trained on the inner training set and evaluated on the inner validation set. The model with the highest AUC score on the inner validation set is then used to calculate the score on the respective test set. Here we report the hyperparameter sets and search strategy that is used for all methods.

#### DeepRC

The set of hyperparameters of DeepRC is shown in Table A3. These hyperparameter combinations are adjusted via a grid search procedure.

**Table A3:** DeepRC hyperparameter search space. Every 5 · 10^3^ updates, the current model was evaluated against the validation fold. The early stopping hyperparameter was determined by selecting the model with the best loss on the validation fold after 10^5^ updates. ^*\**^: Experiments for {64; 128; 256} kernels were omitted for datasets with motif implantation probabilities ≥ 1% in the category “simulated immunosequencing data”.

|  |  |
| --- | --- |
| learning rate | $10^{-4}$ |
| number of kernels ($d_v$) | $\{8; 16; 32; 64^*; 128^*; 256^*\}$ |
| number of CNN layers | $\{1\}$ |
| number of layers in key-NN | $\{2\}$ |
| number of units in key-NN | $\{32\}$ |
| kernel size | $\{5; 7; 9\}$ |
| subsampling sequences | 10,000 |
| batch size | 4 |

#### Known motif

This method does not have hyperparameters and has been applied to all datasets except for the *CMV dataset*.

#### SVM

The corresponding hyperparameter *C* of the SVM is optimized by randomly drawing 10^3^ values in the range of [−6; 6] according to a uniform distribution. These values act as the exponents of a power of 10 and are applied for each of the two kernel types (see Table A4a).

|  |  |
| --- | --- |
| $C$ | $10^{\{-6; 6\}}$ |
| type of kernel | $\{\text{MinMax}; \text{Jaccard}\}$ |
| number of trials | $10^3$ |

Settings used in the hyperparameter search of the SVM baseline approach. The number of trials defines the quantity of random values of the *C* penalty term (per type of kernel).

#### KNN

The amount of neighbors is treated as the hyperparameter and optimized by grid search operating in the discrete range of [1; max {*N*, 10^3^}] with a step size of 1. The corresponding tight upper bound is automatically defined by the total amount of samples *N* ∈ ℕ_*>*0_ in the training set, capped at 10^3^ (see Table A5).

**Table A5:** Settings used in the hyperparameter search of the KNN baseline approach. The number of trials (per type of kernel) is automatically defined by the total amount of samples *N* ∈ ℕ_*>*0_ in the training set, capped at 10^3^.

|  |  |
| --- | --- |
| number of neighbors | $\{1; \max\{N, 10^3\}\}$ |
| type of kernel | $\{\text{MinMax}; \text{Jaccard}\}$ |

#### Logistic regression

The hyperparameter optimization strategy that was used was grid search across hyperparameters given in Table A6.

**Table A6:**
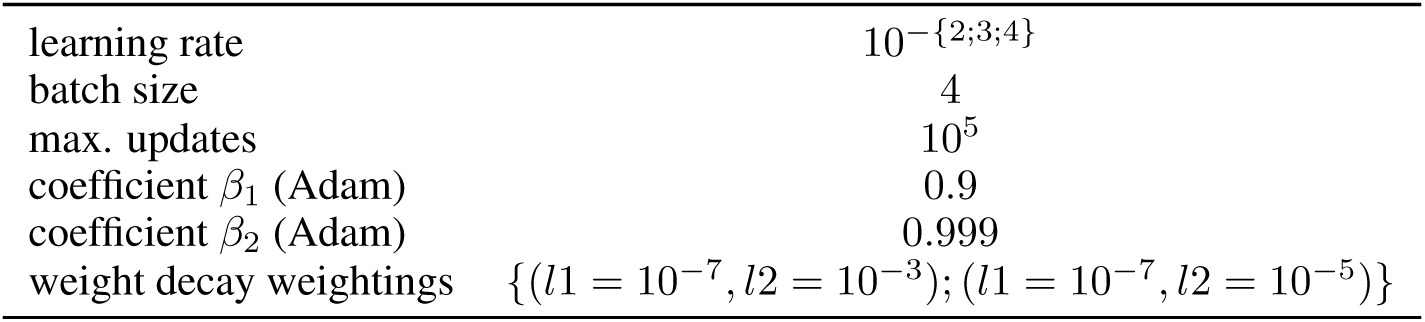
Settings used in the hyperparameter search of the logistic regression baseline approach.

#### Burden test

The burden test selects two hyperparameters: the number of features in the burden set and the type of features, see Table A7.

**Table A7:** Settings used in the hyperparameter search of the burden test approach.

|  |  |
| --- | --- |
| number of features in burden set | $\{50, 100, 150, 250\}$ |
| type of features | $\{4\text{MER}; \text{sequence}\}$ |

#### Logistic MIL

For this method, we adjusted the learning rate as well as the batch size as hyperparameters by randomly drawing 25 different hyperparameter combinations from a uniform distribution. The corresponding range of the learning rate is [−4.5; −1.5], which acts as the exponent of a power of 10. The batch size lies within the range of [1; 32]. For each hyperparameter combination, a model is optimized by gradient descent using Adam, whereas the early stopping parameter is adjusted according to the corresponding validation set (see Table A8).

**Table A8:** Settings used in the hyperparameter search of the logistic MIL baseline approach. The number of trials (per type of relative abundance) defines the quantity of combinations of random values of the learning rate as well as the batch size.

|  |  |
| --- | --- |
| learning rate | $10^{\{-4.5; -1.5\}}$ |
| batch size | $\{1; 32\}$ |
| relative abundance term | $\{4\text{MER}; \text{TCR}\beta\}$ |
| number of trials | 25 |
| max. epochs | $10^2$ |
| coefficient $\beta_1$ (Adam) | 0.9 |
| coefficient $\beta_2$ (Adam) | 0.999 |

### A6 Results

In this section, we report the detailed results on all four categories of datasets (a) simulated immunosequencing data (Table A9) (b) LSTM-generated data (Table A10), (c) real-world data with implanted signals (Table A11), and (d) real-world data on the *CMV dataset* (Table A12), as discussed in the main paper.

**Table A9:** AUC estimates based on 5-fold CV for all 18 datasets in category “simulated immunosequencing data”. The reported errors are standard deviations across the 5 cross-validation folds except for the last column “avg.”, in which they show standard deviations across datasets. Wildcard characters in motifs are indicated by `Z`, characters with 50% probability of being removed by ^d^.

| ID | 0 | 1 | 2 | 3 | 4 | 5 | 6 | 7 | 8 | 9 | 10 | 11 | 12 | 13 | 14 | 15 | 16 | 17 | avg. |
| --- | --- | --- | --- | --- | --- | --- | --- | --- | --- | --- | --- | --- | --- | --- | --- | --- | --- | --- | --- |
| motif freq. $\rho$ | 1% | 0.1% | 0.01% | 1% | 0.1% | 0.01% | 1% | 0.1% | 0.01% | 1% | 0.1% | 0.01% | 1% | 0.1% | 0.01% | 1% | 0.1% | 0.01% | — |
| implanted motif | SFEN | SFEN | SFEN | SF<sup>d</sup>EN | SF<sup>d</sup>EN | SF<sup>d</sup>EN | SFZN | SFZN | SFZN | SF<sup>d</sup>ZN | SF<sup>d</sup>ZN | SF<sup>d</sup>ZN | SZZN | SZZN | SZZN | SZ<sup>d</sup>ZN | SZ<sup>d</sup>ZN | SZ<sup>d</sup>ZN | — |
| DeepRC | **1.000** | **1.000** | 0.703 | **1.000** | **1.000** | 0.600 | **1.000** | **1.000** | 0.509 | **1.000** | **1.000** | 0.492 | **1.000** | **0.997** | 0.487 | 0.999 | **0.942** | 0.492 | **0.864** |
| | $\pm 0.000$ | $\pm 0.000$ | $\pm 0.271$ | $\pm 0.000$ | $\pm 0.000$ | $\pm 0.218$ | $\pm 0.000$ | $\pm 0.000$ | $\pm 0.029$ | $\pm 0.000$ | $\pm 0.001$ | $\pm 0.017$ | $\pm 0.001$ | $\pm 0.002$ | $\pm 0.023$ | $\pm 0.001$ | $\pm 0.048$ | $\pm 0.013$ | $\pm 0.223$ |
| SVM (MinMax) | **1.000** | **1.000** | 0.764 | **1.000** | **1.000** | 0.603 | **1.000** | 0.998 | **0.539** | **1.000** | 0.994 | **0.529** | **1.000** | 0.741 | **0.513** | **1.000** | 0.706 | 0.503 | 0.827 |
| | $\pm 0.000$ | $\pm 0.000$ | $\pm 0.016$ | $\pm 0.000$ | $\pm 0.000$ | $\pm 0.021$ | $\pm 0.000$ | $\pm 0.002$ | $\pm 0.024$ | $\pm 0.000$ | $\pm 0.004$ | $\pm 0.016$ | $\pm 0.000$ | $\pm 0.024$ | $\pm 0.006$ | $\pm 0.000$ | $\pm 0.013$ | $\pm 0.013$ | $\pm 0.210$ |
| SVM (Jaccard) | 0.783 | 0.505 | 0.500 | 0.656 | 0.504 | 0.492 | 0.629 | 0.499 | 0.505 | 0.594 | 0.508 | 0.497 | 0.620 | 0.496 | 0.506 | 0.595 | 0.507 | 0.505 | 0.550 |
| | $\pm 0.010$ | $\pm 0.009$ | $\pm 0.010$ | $\pm 0.009$ | $\pm 0.018$ | $\pm 0.018$ | $\pm 0.011$ | $\pm 0.010$ | $\pm 0.009$ | $\pm 0.007$ | $\pm 0.017$ | $\pm 0.013$ | $\pm 0.007$ | $\pm 0.006$ | $\pm 0.019$ | $\pm 0.013$ | $\pm 0.012$ | $\pm 0.017$ | $\pm 0.080$ |
| KNN (MinMax) | 0.669 | 0.802 | 0.503 | 0.722 | 0.757 | 0.493 | 0.766 | 0.678 | 0.496 | 0.762 | 0.652 | 0.489 | 0.797 | 0.512 | 0.498 | 0.796 | 0.511 | 0.503 | 0.634 |
| | $\pm 0.204$ | $\pm 0.265$ | $\pm 0.038$ | $\pm 0.214$ | $\pm 0.255$ | $\pm 0.017$ | $\pm 0.241$ | $\pm 0.165$ | $\pm 0.014$ | $\pm 0.237$ | $\pm 0.139$ | $\pm 0.015$ | $\pm 0.271$ | $\pm 0.023$ | $\pm 0.014$ | $\pm 0.270$ | $\pm 0.037$ | $\pm 0.006$ | $\pm 0.129$ |
| KNN (Jaccard) | 0.516 | 0.493 | 0.497 | 0.506 | 0.500 | 0.492 | 0.509 | 0.493 | 0.497 | 0.495 | 0.504 | 0.500 | 0.502 | 0.497 | 0.500 | 0.502 | 0.503 | **0.513** | 0.501 |
| | $\pm 0.035$ | $\pm 0.020$ | $\pm 0.013$ | $\pm 0.015$ | $\pm 0.019$ | $\pm 0.014$ | $\pm 0.017$ | $\pm 0.011$ | $\pm 0.018$ | $\pm 0.013$ | $\pm 0.004$ | $\pm 0.017$ | $\pm 0.011$ | $\pm 0.017$ | $\pm 0.022$ | $\pm 0.015$ | $\pm 0.020$ | $\pm 0.012$ | $\pm 0.007$ |
| Logistic regression | **1.000** | **1.000** | **0.786** | **1.000** | **1.000** | **0.607** | **1.000** | 0.997 | 0.527 | **1.000** | 0.992 | 0.526 | **1.000** | 0.719 | 0.505 | **1.000** | 0.694 | 0.510 | 0.826 |
| | $\pm 0.000$ | $\pm 0.000$ | $\pm 0.009$ | $\pm 0.000$ | $\pm 0.000$ | $\pm 0.025$ | $\pm 0.000$ | $\pm 0.002$ | $\pm 0.018$ | $\pm 0.000$ | $\pm 0.004$ | $\pm 0.019$ | $\pm 0.000$ | $\pm 0.019$ | $\pm 0.015$ | $\pm 0.001$ | $\pm 0.021$ | $\pm 0.017$ | $\pm 0.211$ |
| Logistic MIL (KMER) | **1.000** | **1.000** | 0.509 | **1.000** | 0.783 | 0.489 | **1.000** | 0.544 | 0.517 | **1.000** | 0.529 | 0.483 | 0.579 | 0.498 | 0.502 | 0.550 | 0.488 | 0.498 | 0.665 |
| | $\pm 0.000$ | $\pm 0.000$ | $\pm 0.039$ | $\pm 0.000$ | $\pm 0.216$ | $\pm 0.023$ | $\pm 0.000$ | $\pm 0.038$ | $\pm 0.018$ | $\pm 0.000$ | $\pm 0.043$ | $\pm 0.007$ | $\pm 0.042$ | $\pm 0.017$ | $\pm 0.018$ | $\pm 0.051$ | $\pm 0.009$ | $\pm 0.005$ | $\pm 0.224$ |
| Logistic MIL (TCRβ) | 0.544 | 0.505 | 0.493 | 0.487 | 0.476 | 0.500 | 0.520 | 0.495 | 0.510 | 0.492 | 0.506 | 0.503 | 0.509 | 0.505 | 0.500 | 0.475 | 0.489 | 0.500 | 0.501 |
| | $\pm 0.078$ | $\pm 0.014$ | $\pm 0.018$ | $\pm 0.021$ | $\pm 0.019$ | $\pm 0.022$ | $\pm 0.053$ | $\pm 0.009$ | $\pm 0.022$ | $\pm 0.014$ | $\pm 0.019$ | $\pm 0.010$ | $\pm 0.034$ | $\pm 0.009$ | $\pm 0.011$ | $\pm 0.013$ | $\pm 0.024$ | $\pm 0.019$ | $\pm 0.016$ |
| Burden test | 0.770 | 0.523 | 0.510 | 0.666 | 0.510 | 0.509 | 0.652 | 0.508 | 0.505 | 0.583 | 0.508 | 0.509 | 0.564 | 0.508 | 0.507 | 0.536 | 0.508 | 0.504 | 0.549 |
| | $\pm 0.013$ | $\pm 0.013$ | $\pm 0.014$ | $\pm 0.011$ | $\pm 0.009$ | $\pm 0.007$ | $\pm 0.008$ | $\pm 0.011$ | $\pm 0.012$ | $\pm 0.012$ | $\pm 0.007$ | $\pm 0.014$ | $\pm 0.017$ | $\pm 0.010$ | $\pm 0.020$ | $\pm 0.012$ | $\pm 0.016$ | $\pm 0.016$ | $\pm 0.074$ |
| Known motif b. | 1.000 | 1.000 | 0.973 | 1.000 | 1.000 | 0.865 | 1.000 | 1.000 | 0.700 | 1.000 | 0.989 | 0.609 | 1.000 | 0.946 | 0.570 | 1.000 | 0.834 | 0.532 | 0.890 |
| | $\pm 0.000$ | $\pm 0.000$ | $\pm 0.004$ | $\pm 0.000$ | $\pm 0.000$ | $\pm 0.004$ | $\pm 0.000$ | $\pm 0.000$ | $\pm 0.020$ | $\pm 0.000$ | $\pm 0.002$ | $\pm 0.017$ | $\pm 0.000$ | $\pm 0.010$ | $\pm 0.024$ | $\pm 0.000$ | $\pm 0.016$ | $\pm 0.020$ | $\pm 0.168$ |
| Known motif c. | 0.999 | 0.720 | 0.529 | 0.999 | 0.698 | 0.534 | 0.999 | 0.694 | 0.532 | 1.000 | 0.696 | 0.527 | 0.997 | 0.666 | 0.520 | 0.998 | 0.668 | 0.509 | 0.738 |
| | $\pm 0.001$ | $\pm 0.014$ | $\pm 0.020$ | $\pm 0.001$ | $\pm 0.013$ | $\pm 0.017$ | $\pm 0.001$ | $\pm 0.012$ | $\pm 0.012$ | $\pm 0.001$ | $\pm 0.018$ | $\pm 0.018$ | $\pm 0.002$ | $\pm 0.010$ | $\pm 0.009$ | $\pm 0.002$ | $\pm 0.012$ | $\pm 0.013$ | $\pm 0.202$ |

**Table A10:** AUC estimates based on 5-fold CV for all 5 datasets in category “LSTM-generated data”. The reported errors are standard deviations across the 5 cross-validation folds except for the last column “avg.”, in which they show standard deviations across datasets. Characters affected by noise, as described in A3, paragraph “LSTM-generated data”, are indicated by ^r^.

| ID | 0 | 1 | 2 | 3 | 4 | avg. |
| --- | --- | --- | --- | --- | --- | --- |
| motif freq. $\rho$ | 10% | 1% | 0.5% | 0.1% | 0.05% | – |
| implanted motif | $G^r S^r A^r F^r$ | $G^r S^r A^r F^r$ | $G^r S^r A^r F^r$ | $G^r S^r A^r F^r$ | $G^r S^r A^r F^r$ | – |
| DeepRC | **1.000** ± 0.000 | **1.000** ± 0.000 | **1.000** ± 0.000 | **1.000** ± 0.000 | **0.998** ± 0.002 | **1.000** ± 0.001 |
| SVM (MinMax) | **1.000** ± 0.000 | **1.000** ± 0.000 | 0.999 ± 0.001 | 0.999 ± 0.002 | 0.985 ± 0.014 | 0.997 ± 0.007 |
| SVM (Jaccard) | 0.981 ± 0.041 | **1.000** ± 0.000 | **1.000** ± 0.000 | 0.904 ± 0.036 | 0.768 ± 0.068 | 0.931 ± 0.099 |
| KNN (MinMax) | 0.699 ± 0.272 | 0.717 ± 0.263 | 0.732 ± 0.263 | 0.536 ± 0.156 | 0.516 ± 0.153 | 0.640 ± 0.105 |
| KNN (Jaccard) | 0.698 ± 0.285 | 0.606 ± 0.237 | 0.523 ± 0.164 | 0.550 ± 0.186 | 0.539 ± 0.194 | 0.583 ± 0.071 |
| Logistic regression | **1.000** ± 0.000 | **1.000** ± 0.000 | 0.934 ± 0.147 | 0.604 ± 0.193 | 0.427 ± 0.156 | 0.793 ± 0.262 |
| Logistic MIL (KMER) | 0.997 ± 0.004 | 0.718 ± 0.112 | 0.637 ± 0.144 | 0.571 ± 0.146 | 0.528 ± 0.129 | 0.690 ± 0.186 |
| Logistic MIL (TCRβ) | 0.541 ± 0.086 | 0.566 ± 0.162 | 0.468 ± 0.086 | 0.505 ± 0.067 | 0.500 ± 0.121 | 0.516 ± 0.038 |
| Burden test | **1.000** ± 0.000 | **1.000** ± 0.000 | **1.000** ± 0.000 | 0.999 ± 0.003 | 0.792 ± 0.280 | 0.958 ± 0.093 |
| Known motif b. | 1.000 ± 0.000 | 1.000 ± 0.000 | 1.000 ± 0.000 | 0.999 ± 0.003 | 0.999 ± 0.003 | 1.000 ± 0.001 |
| Known motif c. | 1.000 ± 0.000 | 1.000 ± 0.000 | 0.989 ± 0.011 | 0.722 ± 0.085 | 0.626 ± 0.094 | 0.867 ± 0.180 |

**Table A11:** AUC estimates based on 5-fold CV for all 4 datasets in category “real-world data with implanted signals”. The reported errors are standard deviations across the 5 cross-validation folds except for the last column “avg.”, in which they show standard deviations across datasets. **OM 1%:** In this dataset, a single motif with a frequency of 1% was implanted. **OM 0.1%:** In this dataset, a single motif with a frequency of 0.1% was implanted. **MM 1%:** In this dataset, multiple motifs with a frequency of 1% were implanted. **MM 0.1%:** In this dataset, multiple motifs with a frequency of 0.1% were implanted. A detailed description of the motifs is provided in Section A3, paragraph “Real-world data with implanted signals.”.

|  | OM 1% | OM 0.1% | MM 1% | MM 0.1% | Avg. |
| --- | --- | --- | --- | --- | --- |
| DeepRC | **1.000** ± 0.000 | **0.984** ± 0.008 | 0.999 ± 0.001 | **0.938** ± 0.009 | **0.980** ± 0.029 |
| SVM (MinMax) | **1.000** ± 0.000 | 0.578 ± 0.020 | **1.000** ± 0.000 | 0.531 ± 0.019 | 0.777 ± 0.258 |
| SVM (Jaccard) | 0.988 ± 0.003 | 0.527 ± 0.016 | **1.000** ± 0.000 | 0.574 ± 0.019 | 0.772 ± 0.257 |
| KNN (MinMax) | 0.744 ± 0.237 | 0.486 ± 0.031 | 0.674 ± 0.182 | 0.500 ± 0.022 | 0.601 ± 0.128 |
| KNN (Jaccard) | 0.652 ± 0.155 | 0.484 ± 0.025 | 0.695 ± 0.200 | 0.508 ± 0.025 | 0.585 ± 0.104 |
| Logistic regression | **1.000** ± 0.000 | 0.544 ± 0.035 | 0.991 ± 0.003 | 0.512 ± 0.035 | 0.762 ± 0.270 |
| Logistic MIL (KMER) | 0.541 ± 0.074 | 0.506 ± 0.034 | 0.994 ± 0.004 | 0.620 ± 0.153 | 0.665 ± 0.224 |
| Logistic MIL (TCRβ) | 0.503 ± 0.032 | 0.501 ± 0.016 | 0.992 ± 0.003 | 0.782 ± 0.030 | 0.695 ± 0.238 |
| Burden test | **1.000** ± 0.000 | 0.640 ± 0.048 | **1.000** ± 0.000 | 0.891 ± 0.016 | 0.883 ± 0.170 |
| Known motif b. | 1.000 ± 0.000 | 0.704 ± 0.028 | 0.994 ± 0.003 | 0.620 ± 0.038 | 0.830 ± 0.196 |
| Known motif c. | 0.920 ± 0.004 | 0.562 ± 0.028 | 0.647 ± 0.030 | 0.515 ± 0.031 | 0.661 ± 0.181 |

**Table A12:** Results on the *CMV dataset* (real-world data) in terms of AUC, F1 score, balanced accuracy, and accuracy. For F1 score, balanced accuracy, and accuracy, all methods use their default thresholds. Each entry shows mean and standard deviation across 5 cross-validation folds.

|  | AUC | F1 score | Balanced accuracy | Accuracy |
| --- | --- | --- | --- | --- |
| DeepRC | **0.831** ± 0.002 | **0.726** ± 0.050 | **0.741** ± 0.043 | 0.727 ± 0.049 |
| SVM (MinMax) | 0.825 ± 0.022 | 0.680 ± 0.056 | 0.734 ± 0.037 | **0.742** ± 0.031 |
| SVM (Jaccard) | 0.546 ± 0.021 | 0.272 ± 0.184 | 0.523 ± 0.026 | 0.542 ± 0.032 |
| KNN (MinMax) | 0.679 ± 0.076 | 0.000 ± 0.000 | 0.500 ± 0.000 | 0.545 ± 0.044 |
| KNN (Jaccard) | 0.534 ± 0.039 | 0.073 ± 0.101 | 0.508 ± 0.012 | 0.551 ± 0.042 |
| Logistic regression | 0.607 ± 0.058 | 0.244 ± 0.206 | 0.552 ± 0.049 | 0.590 ± 0.019 |
| Logistic MIL (KMER) | 0.582 ± 0.065 | 0.118 ± 0.264 | 0.503 ± 0.007 | 0.515 ± 0.058 |
| Logistic MIL (TCRβ) | 0.515 ± 0.073 | 0.000 ± 0.000 | 0.496 ± 0.008 | 0.541 ± 0.039 |
| Burden test | 0.699 ± 0.041 | - | - | - |

### A7 Repertoire generation via LSTM

We trained a conventional next-character LSTM model (Graves, 2013) based on the implementation in https://github.com/spro/practical-pytorch (access date 1st of May, 2020) using `PyTorch 1.3.1` (Paszke et al., 2019). For this, we applied an LSTM model with 100 LSTM blocks in 2 layers, which was trained for 5, 000 epochs using the Adam optimizer (Kingma & Ba, 2014) with learning rate 0.01, an input batch size of 100 character chunks, and a character chunk length of 200. As input we used the immuno-sequences in the `CDR3` column of the *CMV dataset*, where we repeated sequences according to their counts in the repertoires, as specified in the `templates` column of the *CMV dataset*. We excluded repertoires with unknown CMV status and unknown sequence abundance from training.

After training, we generated 1, 000 repertoires using a `temperature` value of 0.8. The number of sequences per repertoire was sampled from a Gaussian 𝒩 (*µ* = 285*k, σ* = 156*k*) distribution, where the whole repertoire was generated by the LSTM at once. That is, the LSTM can base the generation of the individual AA sequences in a repertoire, including the AAs and the lengths of the sequences, on the generated repertoire. A random immuno-sequence from the trained-on repertoires was used as initialization for the generation process. This immuno-sequence was not included in the generated repertoire.

Finally, we randomly assigned 500 of the generated repertoires to the positive (diseased) and 500 to the negative (healthy) class. We then implanted motifs in the positive class repertoires as described in Section A3.2.

As illustrated in the comparison of histograms given in Fig. A2, the generated immuno-sequences exhibit a very similar distribution of 4-mers and AAs compared to the original *CMV dataset*.

**Figure A2:**
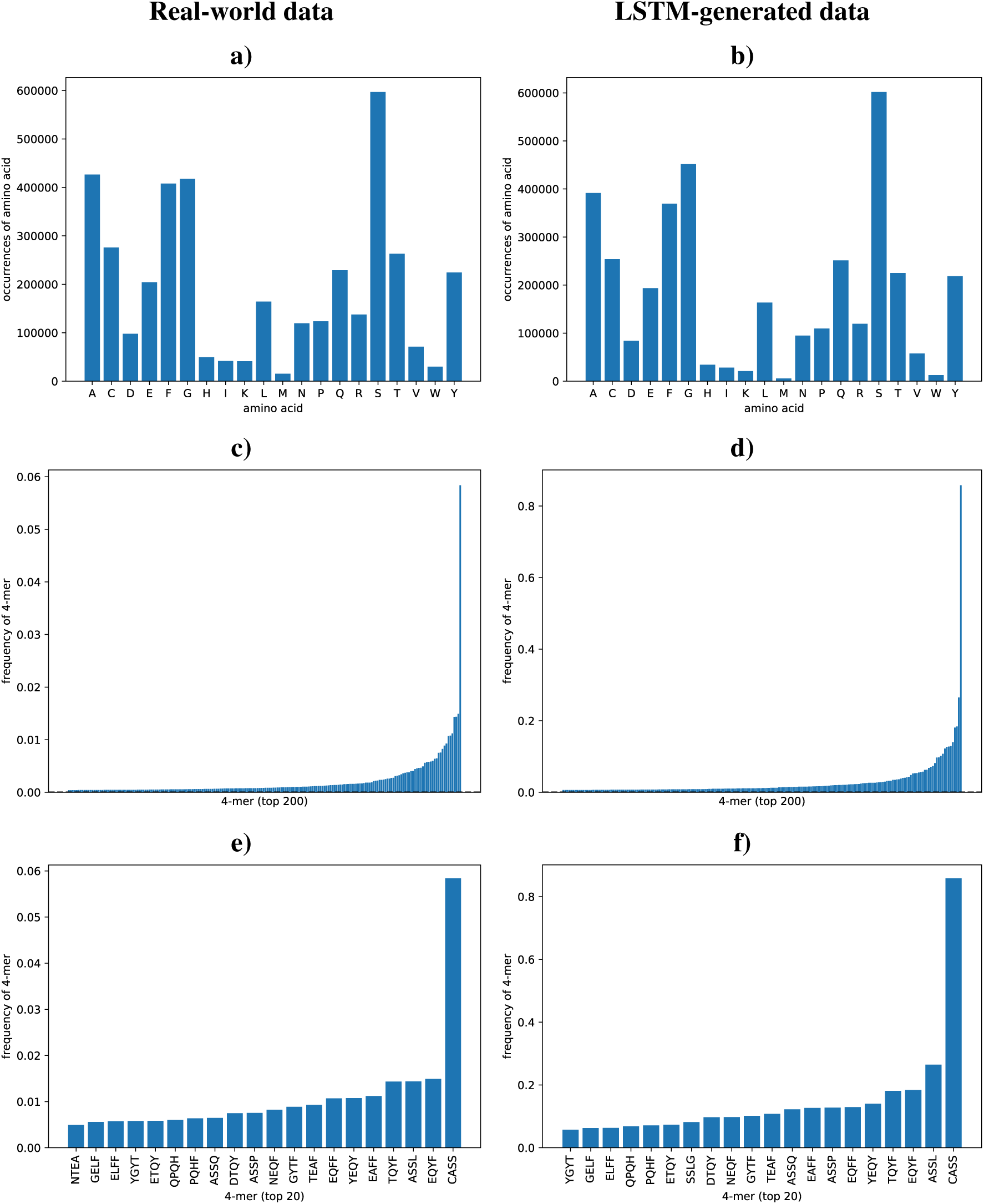
Distribution of AAs and k-mers in real-world *CMV dataset* and LSTM-generated data. **Left:** Histograms of real-world data. **Right:** Histograms of LSTM-generated data. **a)** Frequency of AAs in sequences of the *CMV dataset*. **b)** Frequency of AAs in sequences of the LSTM-generated datasets. **c)** Frequency of top 200 4-mers in sequences of the *CMV dataset*. **d)** Frequency of top 200 4-mers in sequences of the LSTM-generated datasets. **e)** Frequency of top 20 4-mers in sequences of the *CMV dataset*. **f)** Frequency of top 20 4-mers in sequences of the LSTM-generated datasets. Overall the distributions of AAs and 4-mers are similar in both datasets.

#### A8 Interpreting DeepRC

DeepRC allows for two forms of interpretability methods. (a) Due to its attention-based design, a trained model can be used to compute the attention weights of a sequence, which directly indicates its importance. (b) DeepRC furthermore allows for the usage of contribution analysis methods, such as Integrated Gradients (IG) (Sundararajan et al., 2017) or Layer-Wise Relevance Propagation (Montavon et al., 2018; Arras et al., 2019; Montavon et al., 2019; Preuer et al., 2019). We apply IG to identify the input patterns that are relevant for the classification. To identify AA patterns with high contributions in the input sequences, we apply IG to the AAs in the input sequences. Additionally, we apply IG to the kernels of the 1D CNN, which allows us to identify AA motifs with high contributions. In detail, we compute the IG contributions for the AAs and positional features in the kernels for every repertoire in the validation and test set, so as to exclude potential artifacts caused by over-fitting. Averaging the IG values over these repertoires then results in concise AA motifs. We include qualitative visual analyses of the IG method on different datasets below.

Here, we provide examples for the interpretation of trained DeepRC models using Integrated Gradients (IG) (Sundararajan et al., 2017) as contribution analysis method. The following illustrations were created using 50 IG steps, which we found sufficient to achieve stable IG results.

A visual analysis of DeepRC models on the simulated datasets, as illustrated in Tab. A13 and Fig. A3, shows that the implanted motifs can be successfully extracted from the trained model and are straight-forward to interpret. In the real-world *CMV dataset*, DeepRC finds complex patterns with high variability in the center regions of the immuno-sequences, as illustrated in figure A4.

**Table A13:**
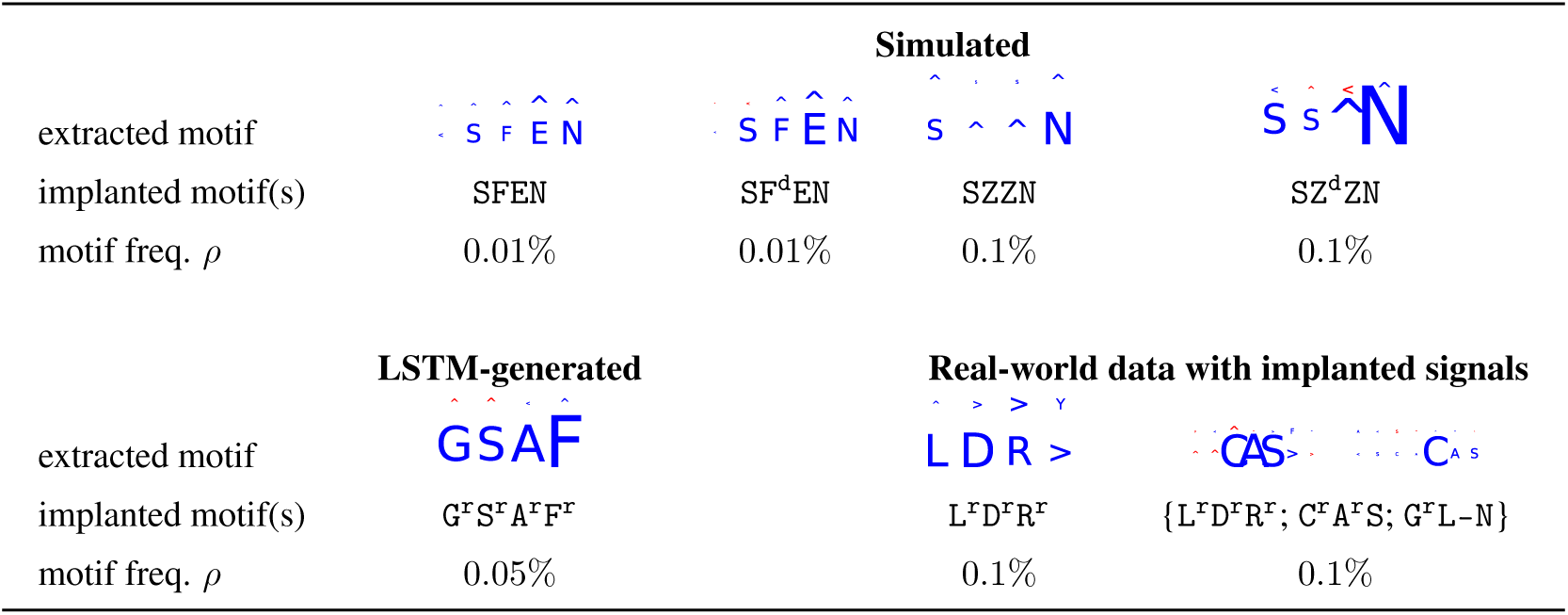
Visualization of motifs extracted from trained DeepRC models for datasets from categories “simulated immunosequencing data”, “LSTM-generated data”, and “real-world data with implanted signals”. Motif extraction was performed using Integrated Gradients on the 1D CNN kernels over the validation set and test set repertoires of one CV fold. Wildcard characters are indicated by Z, random noise on characters by ^r^, characters with 50% probability of being removed by ^d^, and gap locations of random lengths of {0; 1; 2} by -. Larger characters in the extracted motifs indicate higher contribution, with blue indicating positive contribution and red indicating negative contribution towards the prediction of the diseased class. Contributions to positional encoding are indicated by *<* (beginning of sequence), ∧ (center of sequence), and *>* (end of sequence). Only kernels with relatively high contributions are shown, i.e. with contributions roughly greater than the average contribution of all kernels.

**Figure A3:**
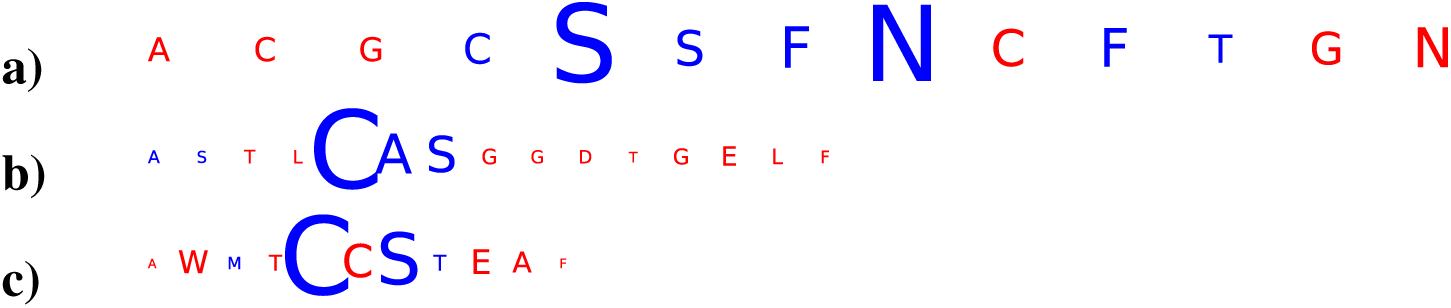
Integrated Gradients applied to input sequences of positive class repertoires. Three sequences with the highest contributions to the prediction of their respective repertoires are shown. **a)** Input sequence taken from “simulated immunosequencing data” with implanted motif SZ^d^Z^d^N and motif implantation probability 0.1%. The DeepRC model reacts to the `S` and `N` at the 5^th^ and 8^th^ sequence position, thereby identifying the implanted motif in this sequence. **b)** and **c)** Input sequence taken from “real-world data with implanted signals” with implanted motifs {L^r^D^r^R^r^; C^r^A^r^S; G^r^L-N} and motif implantation probability 0.1%. The DeepRC model reacts to the fully implanted motif `CAS` (b) and to the partly implanted motif `AAs C` and `A` at the 5^th^ and 7^th^ sequence position (c), thereby identifying the implanted motif in the sequences. Wildcard characters in implanted motifs are indicated by `Z`, characters with 50% probability of being removed by ^d^, and gap locations of random lengths of 0; 1; 2 by -. Larger characters in the sequences indicate higher contribution, with blue indicating positive contribution and red indicating negative contribution towards the prediction of the diseased class.

**Figure A4:**
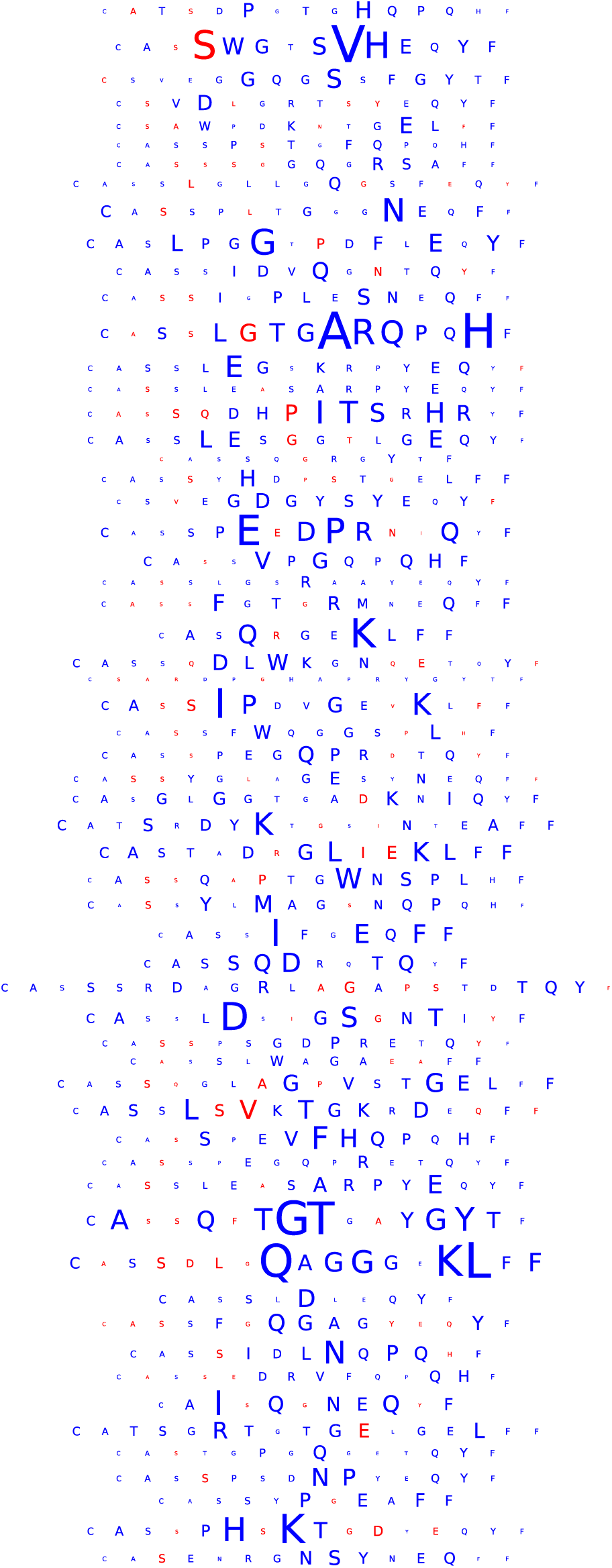
Visualization of the contributions of characters within a sequence via IG. Each sequence was selected from a different repertoire and showed the highest contribution in its repertoire. The model was trained on *CMV dataset*, using a kernel size of 9, 32 kernels and 137 repertoires for early stopping. Larger characters in the extracted motifs indicate higher contribution, with blue indicating positive contribution and red indicating negative contribution towards the prediction of the disease class.

### A9 Attention values for previously associated CMV sequences

Table A14 lists sequences of the *CMV dataset* that were previously associated with CMV immune status and their assigned high attention values by DeepRC.

**Table A14:**
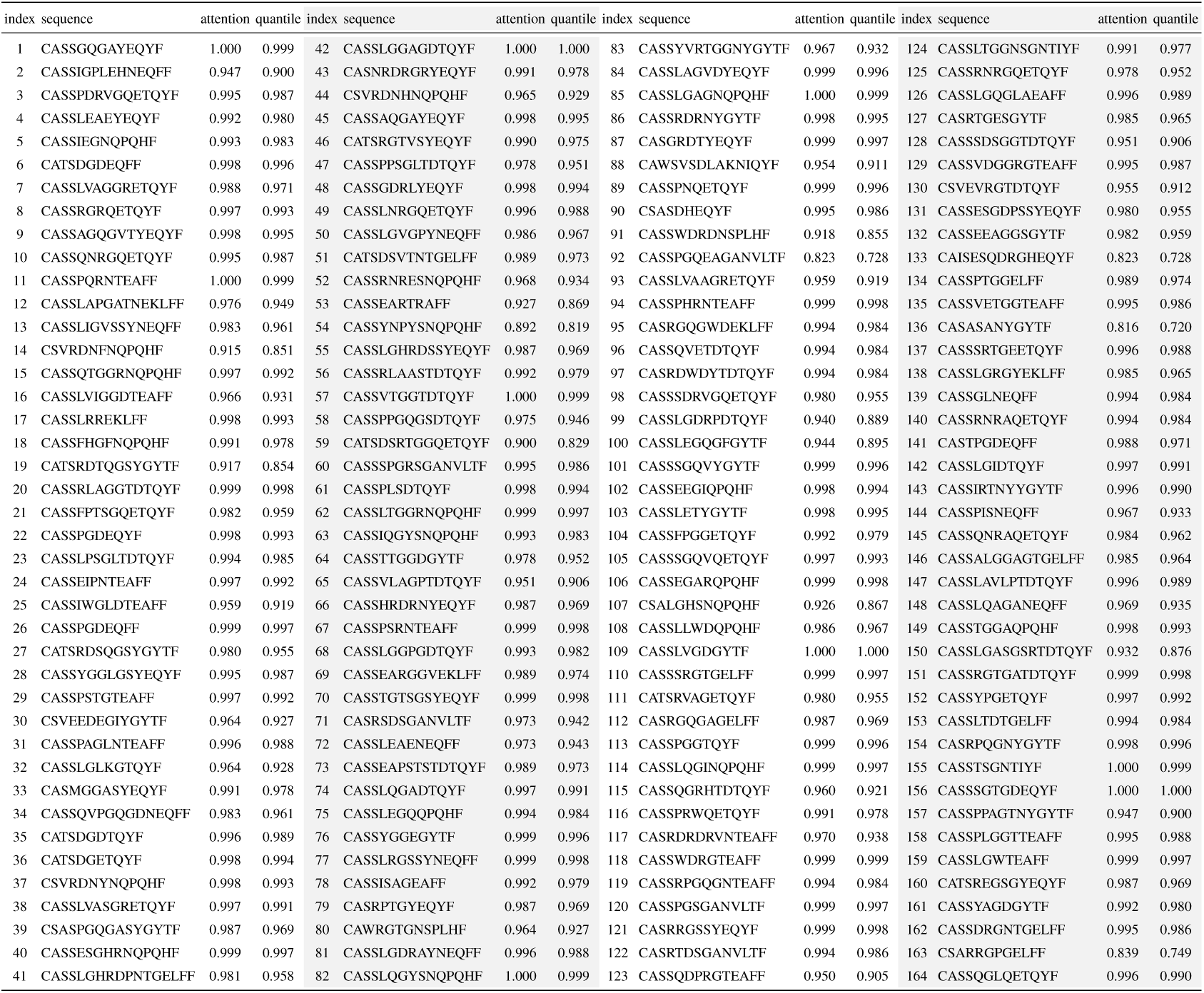
TCRβ sequences that had been discovered by Emerson et al. (2017) with their associated attention values by DeepRC. These sequences have significantly (*p*-value 1.3e-93) higher attention values than other sequences. The column “quantile” provides the quantile values of the empiricial distribution of attention values across all sequences in the dataset.

### A10 DeepRC variations and ablation study

In this section we investigate the impact of different variations of DeepRC on the performance on the *CMV dataset*. We consider both a CNN-based sequence embedding, as used in the main paper, and an LSTM-based sequence embedding. In both cases we vary the number of attention heads and the *β* parameter for the softmax function the attention mechanism (see Eq. 2 in main paper). For the CNN-based sequence embedding we also vary the number of CNN kernels and the kernel sizes used in the 1D CNN. For the LSTM-based sequence embedding we use one one-directional LSTM layer, of which the output values at the last sequence position (without padding) are taken as embedding of the sequence. Here we vary the number of LSTM blocks in the LSTM layer. To counter over-fitting due to the increased complexity of these DeepRC variations, we added a *l*2 weight penalty to the training loss. The factor with which the *l*2 weight penalty contributes to the training loss is varied over 3 orders of magnitudes, where suitable value ranges were manually determined on one of the training folds beforehand.

To reduce the computational effort, we do not consider all numbers of kernels that were considered in the main paper. Furthermore, we only compute the AUC scores on 3 of the 5 cross-validation folds. The hyperparameters, which were used in a grid search procedure, are listed in Tab. A15 for the CNN-based sequence embedding and Tab. A16 for the LSTM-based sequence embedding.

#### Results

We show performance in terms of AUC score with single hyperparameters set to fixed values so as to investigate their influence in Tab. A18 for the CNN-based sequence embedding and Tab. A17 for the LSTM-based sequence embedding. We note that due to restricted computational resources this study was conducted with fewer different numbers of CNN kernels, with the AUC estimated from only 3 of the 5 cross-validation folds, which leads to a slight decrease of performance in comparison to the full hyperparameter search and cross-validation procedure used in the main paper. As can be seen in Tab. A18 and A17, the LSTM-based sequence embedding generalizes slightly better than the CNN-based sequence embedding. The performance of DeepRC, however, remains rather robust w.r.t. the different hyperparameter settings.

**Table A15:** Hyperparameter search space for DeepRC variations with CNN-based sequence embedding. Every 5 ° 10^3^ updates, the current model was evaluated against the validation fold. The early stopping hyperparameter was determined by selecting the model with the best loss on the validation fold after 2 · 10^5^ updates.

|  |  |
| --- | --- |
| learning rate | $10^{-4}$ |
| number of attention heads | {1; 16; 64} |
| $\beta$ of attention softmax | {0.1; 1.0; 10.0} |
| $l_2$ weight penalty | {1.0; 0.1; 0.01} |
| number of kernels | {8; 32; 128} |
| number of CNN layers | {1} |
| number of layers in key-NN | {2} |
| number of units in key-NN | {32} |
| kernel size | {5; 7; 9} |
| subsampling sequences | 10,000 |
| batch size | 4 |

**Table A16:** Hyperparameter search space for DeepRC variations with LSTM-based sequence embedding. Every 5 · 10^3^ updates, the current model was evaluated against the validation fold. The early stopping hyperparameter was determined by selecting the model with the best loss on the validation fold after 2 · 10^5^ updates.

|  |  |
| --- | --- |
| learning rate | $10^{-4}$ |
| number of attention heads | {1; 16; 64} |
| $\beta$ of attention softmax | {0.1; 1.0; 10.0} |
| $l_2$ weight penalty | {0.01; 0.001; 0.0001} |
| number of LSTM blocks | {8; 32; 128} |
| number of CNN layers | {1} |
| number of layers in key-NN | {2} |
| number of units in key-NN | {32} |
| subsamped sequences | 10,000 |
| batch size | 4 |

**Table A17:**
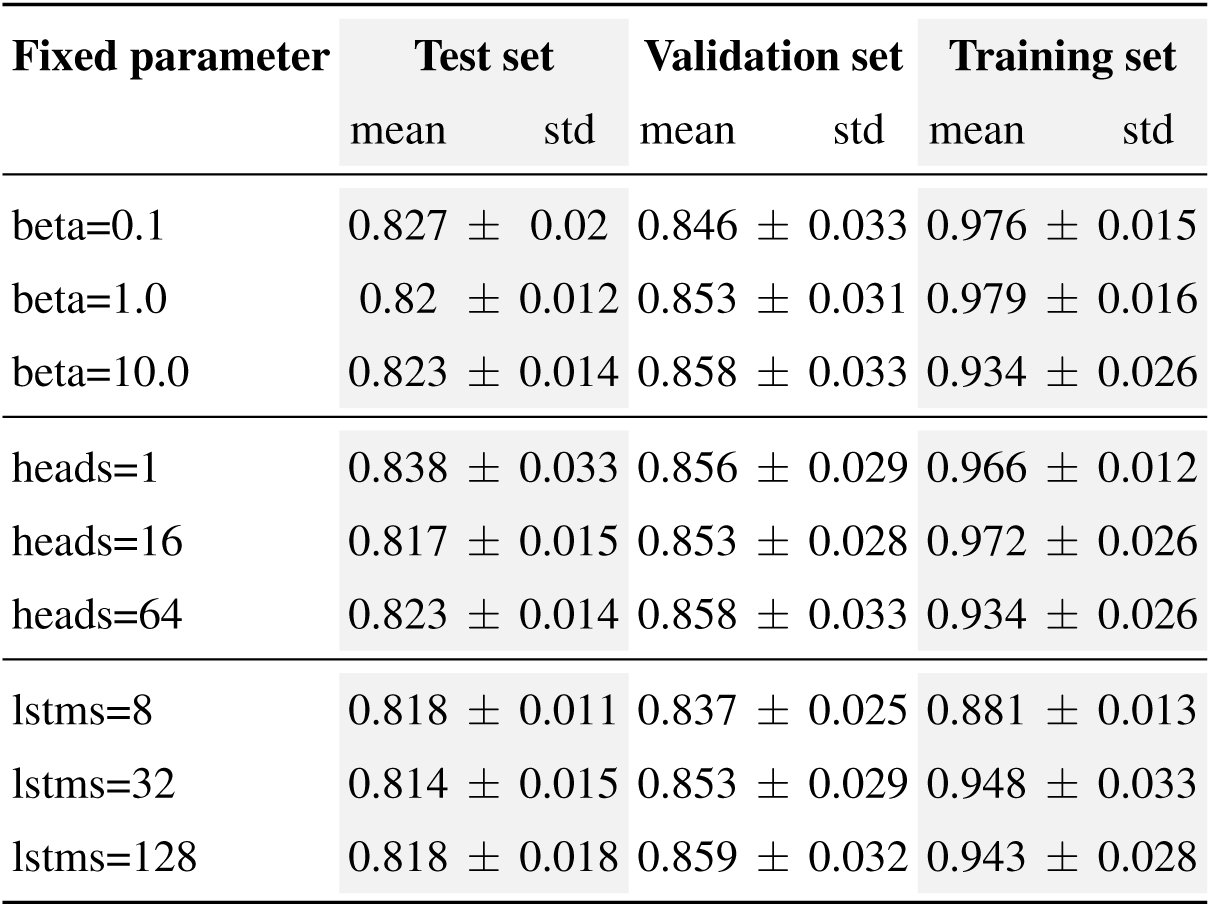
Impact of hyperparameters on DeepRC with LSTM for sequence encoding. Mean (“mean”) and standard deviation (“std”) for the area under the ROC curve over the first 3 folds of a 5-fold nested cross-validation for different sub-sets of hyperparameters (“sub-set”) are shown. The following sub-sets were considered: “full”: Full grid search over hyperparameters; “beta=*”: Grid search over hyperparameters with reduction to specific value * of beta value of attention softmax; “heads=*”: Grid search over hyperparameters with reduction to specific number * of attention heads; “lstms=*”: Grid search over hyperparameters with reduction to specific number * of LSTM blocks for sequence embedding.

**Table A18:**
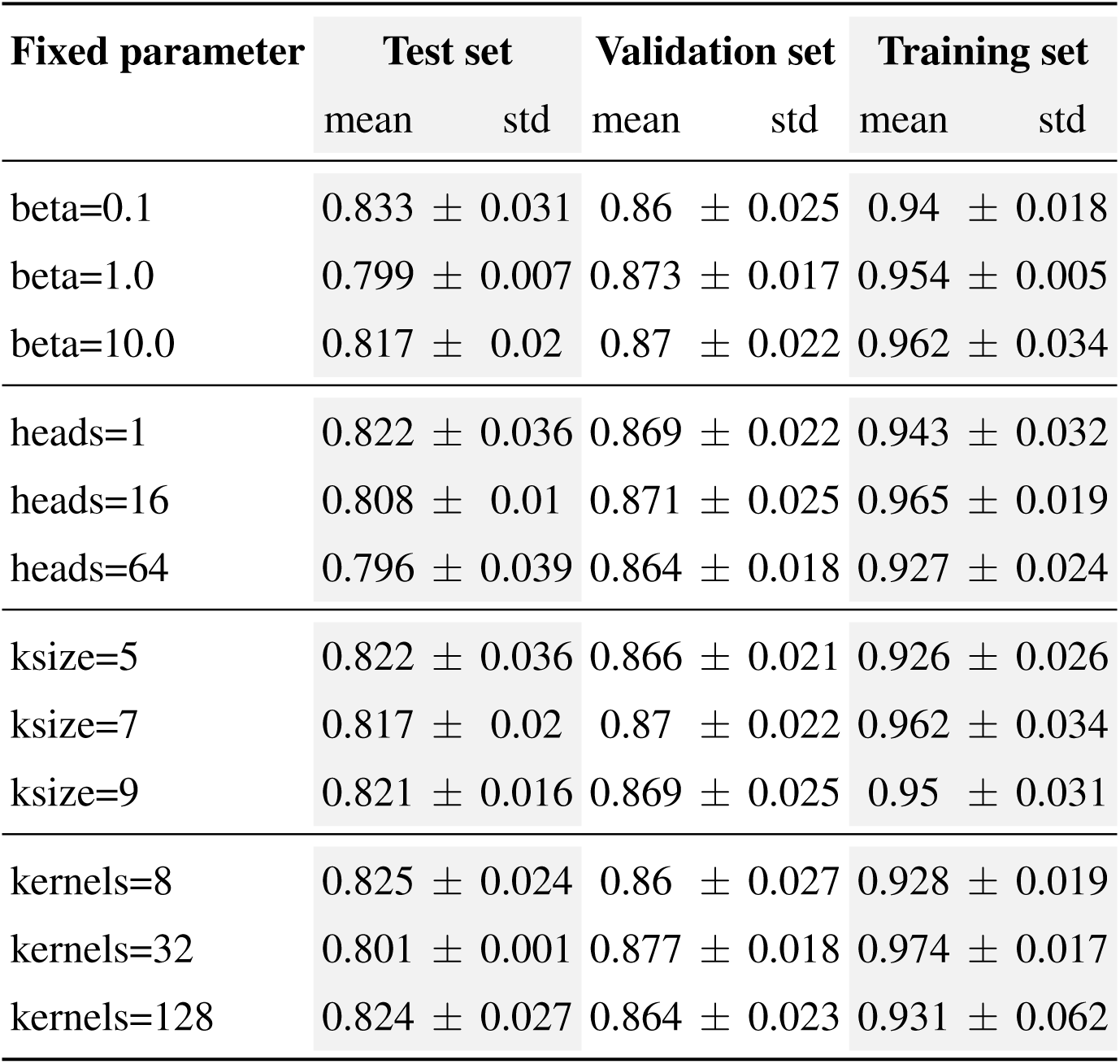
Impact of hyperparameters on DeepRC with 1D CNN for sequence encoding. Mean (“mean”) and standard deviation (“std”) for the area under the ROC curve over the first 3 folds of a 5-fold nested cross-validation for different sub-sets of hyperparameters (“sub-set”) are shown. The following sub-sets were considered: “full”: Full grid search over hyperparameters; “beta=*”: Grid search over hyperparameters with reduction to specific value * of beta value of attention softmax; “heads=*”: Grid search over hyperparameters with reduction to specific number * of attention heads; “ksize=*”: Grid search over hyperparameters with reduction to specific kernel size * of 1D CNN kernels for sequence embedding; “kernels=*”: Grid search over hyperparameters with reduction to specific number * of 1D CNN kernels for sequence embedding.

